# SMG7 and eIF4A constitute a homeostatic module controlling P-body condensation and function of Meiotic bodies

**DOI:** 10.1101/2025.05.03.652043

**Authors:** Albert Cairo, Neha Shukla, Sofia Kanavorova, Jan Skalak, Pavlina Mikulkova, Anna Vargova, David Potesil, Zbyněk Zdráhal, Jan Hejatko, Karel Riha

## Abstract

Processing bodies (P-bodies) are ribonucleoprotein condensates that regulate RNA processing and storage. Although present constitutively in most cells, their size and composition dynamically change in response to developmental and environmental cues. However, mechanisms governing P-body assembly and remodelling remain poorly understood. Here we show that in *Arabidopsis*, SMG7 interacts with the eIF4A helicases and recruits them to P-bodies. eIF4As limit P-bodies condensation and also restrict stress granule (SG) formation under heat stress. We further identify meiotic (M-)bodies as composite RNP granules with a P-body core surrounded by a SG-like shell. The SMG7-eIF4A module regulates the recruitment of the meiosis-specific protein TDM1 into M-bodies, influencing meiotic exit and plant reproduction. Our findings suggest that SMG7 functions as an adaptor protein that recruits client proteins into P-bodies and, together with eIF4A, forms a regulatory module that regulates P-body composition and maintains their size homeostasis.

## Introduction

Ribonucleoprotein (RNP) granules, such as stress granules (SGs), P-bodies or Cajal bodies, are assemblies of proteins and RNAs that are either stably present in cells, or form in response to stress or developmental stimuli. These membrane-less organelles serve as dynamic hubs co-ordinating various steps of RNA biogenesis and processing, including transcription, splicing, storage, or translational repression, ensuring tight control over gene expression. RNP granules are held together by multivalent low-affinity interactions allowing constant exchange of proteins and RNAs with their surroundings, which contributes to their dynamic nature (1, 2).

RNAs are not merely passive subjects of RNP granule regulation, but act as important structural scaffold for their nucleation and assembly. As long flexible polymers, RNAs offer numerous sites for intermolecular interactions with either other RNAs or RNA-binding proteins, which can through binding to other proteins connect different RNPs. These multivalent interactions facilitate the formation of higher order assemblies (3). Notably, RNA exhibits an intrinsic propensity for self-assembly at relatively low concentrations *in vitro*, and this characteristic has been implicated in the nucleation of stress granules that consist of translationally-repressed mRNAs released from ribosomes during cellular stress (4–7). Because the intracellular concentration of RNA surpasses the *in vitro* threshold for self-assembly, exposed RNAs can engage in promiscuous interactions that drive the spontaneous formation of SGs and other RNP condensates.

Uncontrolled RNA self-assembly can disrupt cellular homeostasis by sequestering essential RNA binding proteins, interfering with RNA processing, or promoting the formation of aberrant condensates. Recent studies in yeast and human cells have suggested the existence of an RNA chaperone network that counteracts the formation of these assemblies, facilitates RNP remodelling and regulates RNP granule homeostasis (6, 8–11). These processes can regulate RNA flux between condensates and the surrounding cytoplasm, thereby influencing mRNA availability and translatability (11, 12). One example of RNA chaperone is eIF4A, which is primarily known for its role in the initiation of translation. eIF4A is the archetypal member of the DEAD-box helicase family and, together with the scaffold protein eIF4G and the mRNA 5’ cap-binding protein eIF4E, forms the eukaryotic translation initiation complex eIF4F (13, 14). eIF4A functions by unwinding structures within 5’-UTR of mRNAs, hereby facilitating the scanning for start codons by the 43S-preinitiation ribosome complex (15). Separate from its function in translation, eIF4A has also been shown to limit RNA condensation and SG formation in human cells (6).

P-bodies are another type of RNP condensates related to SGs that accumulates translationally repressed mRNAs. However, unlike SGs, P-bodies are typically constitutively present in cells and are enriched in proteins involved in mRNA degradation, such as non-sense mediated RNA decay (NMD) factors, decapping enzymes, exonucleases, and various RNA-binding proteins, which collectively regulate mRNA turnover and storage (16, 17). In plants, P-bodies have been implicated in a number of biological processes, including pathogen defence, photomorphogenesis, actin remodelling, heat stress adaptation, and meiotic progression (18–22). Although present in most plant cells, P-bodies are highly dynamic structures, and their size and compositions change in response to environmental and developmental stimuli (18, 22–24). This suggests the existence of active remodelling mechanisms that regulate P-body assembly, disassembly, and compositional plasticity, allowing cells to fine-tune mRNA metabolism according to changing physiological requirements.

SMG7 is an evolutionary conserved NMD factor and a core component of P-bodies in both mammals and plants (20, 25). In NMD, SMG7 binds to phosphorylated UPF1-helicase through its 14-3-3 like domain and recruits it to P-bodies (26–28). In *Arabidopsis*, SMG7 also fulfils an NMD-independent function in meiotic exit (29, 30). At the end of meiosis, SMG7 binds and recruits the meiosis specific protein TDM1 into P-bodies. TDM1 interacts with the translation initiation factor eIFiso4G2 and sequesters it in P-bodies, thereby inhibiting translation and facilitating meiotic exit (22). Thus, during meiosis SMG7 changes P-body composition and alters its function to temporarily inhibit translation, enabling meiotic exit.

14-3-3 proteins are known to interact with a broad spectrum of substates (31), raising the possibility that SMG7, which contains a 14-3-3-like domain, may associate with additional proteins beyond TDM1 and UPF1. In this study, we identify *Arabidopsis* DEAD-box helicases eIF4A1 and eIF4A2 as novel SMG7 interactors and uncover a previously unrecognized role for SMG7 in P-body remodelling. Similar to its interactions with UPF1 and TDM1, SMG7 binds with eIF4A1/2 and recruits them to meiotic P-bodies. We further show that meiotic cells contain specialized cytoplasmic RNP condensates, M-bodies, that consist of P-body core surrounded by a shell enriched with SG components. Consistent with their RNA chaperone activity, eIF4A1/2 limit the size of both SGs and P-bodies in somatic cells, as well as the size of M-bodies, influencing their capacity to recruit TDM1 and to promote meiotic exit. We propose that the SMG7-eIF4A constitutes an autoregulatory module controlling P-body remodelling and homeostasis.

## Results

### SMG7 interacts with the RNA helicase eIF4A

To identify SMG7 interactors, we used the *Arabidopsis* line expressing SMG7-MYC (Figure 1A), which complements the *smg7-1* null mutation (30, 32), for immunoprecipitation (Figure 1B) followed by mass spectrometry. Among the most enriched proteins were the two isoforms of eIF4A, eIF4A1 and eIF4A2 (Table S1). The *Arabidopsis* genome encodes three eIF4A orthologues. The cytoplasmic eIF4A1 and eIF4A2 act in translation and exhibit 97% sequence similarity and similar expression patterns (33), whereas eIF4A3 is part of the exon-junction complex and shuttles between nucleus and cytoplasm (34). While the *eif4a1* and *eif4a2* single mutants are viable, showing no phenotype, or in the case of *eif4a1* slightly impaired growth and fertility, deficiency in both genes is lethal, indicating a substantial functional redundancy (33). To validate the interaction, we generated the *eIF4A1:YFP* line containing the genic region of *eIF4A1* fused with YFP that complemented the *eif4a1* phenotype, and crossed it with the *SMG7-MYC* line. Co-immunoprecipitation of SMG7-MYC with the GFP nanobeads from plants expressing both constructs indicate that SMG7 and eIF4A1 indeed interact (Figure 1C).

**Figure 1.**
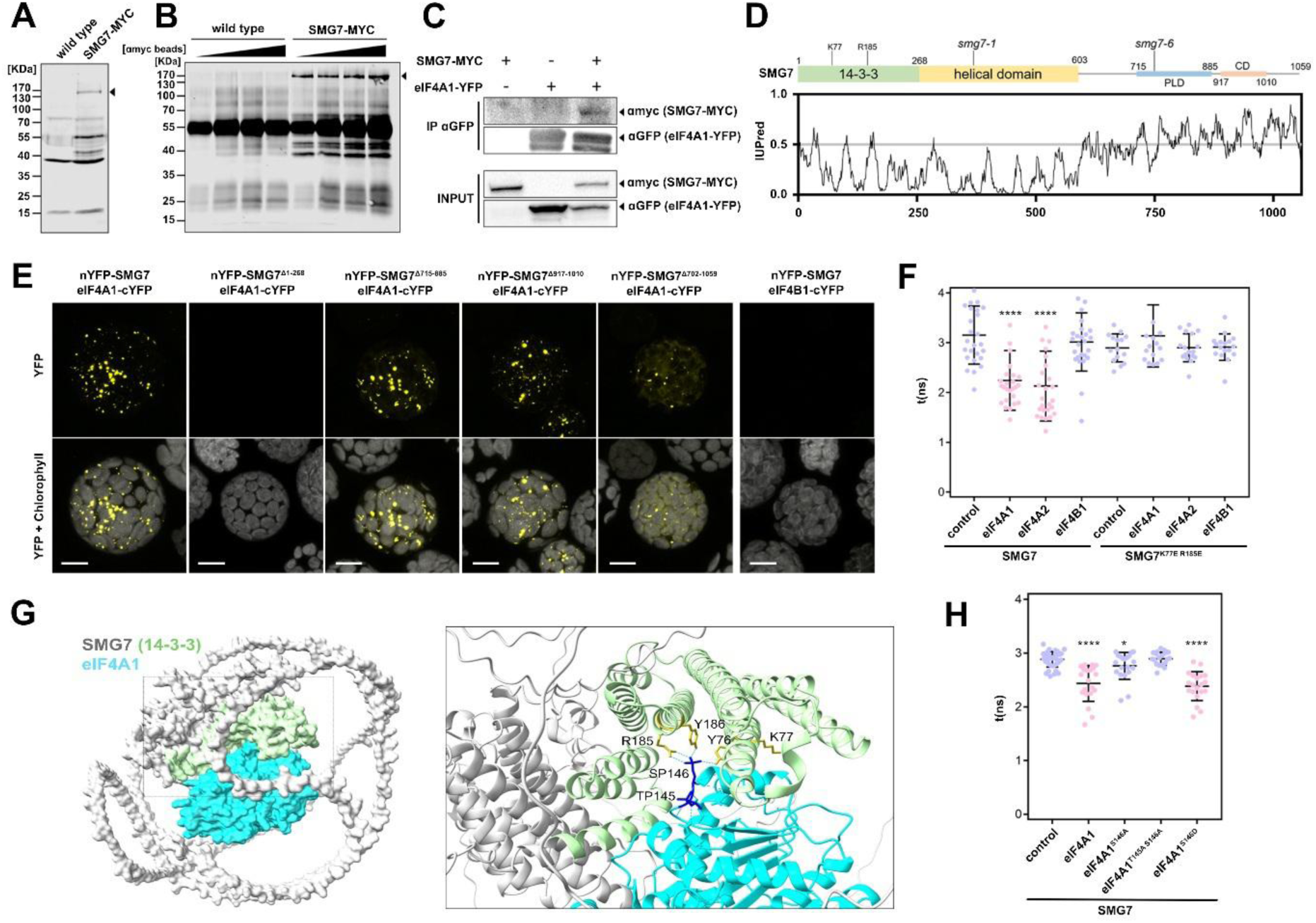
SMG7 interacts with the RNA helicase eIF4A in P-bodies. **(A)** Western blot analysis of SMG7-MYC expression in *SMG7:MYC* plants. **(B)** Western blot analysis of immunoprecipitates of protein extracts of wild type and *SMG7:MYC* plants using increasing amount of anti-MYC antibody. **(C)** Co-immunoprecipitation of SMG7-MYC and eIF4A-YFP from plants expressing either *SMG7-MYC*, or *eIF4A1-YFP* or both constructs using anti-GFP antibody. SMG7-MYC was detected with anti-MYC antibody and eIF4A1 with anti-GFP antibody. **(D)** SMG7 domain structure indicating the T-DNA insertion site of the mutants *smg7-1* and *smg7-6*. Prediction of disordered regions using IUPred (76) is depicted bellow. **(E)** BiFC assay in *Arabidopsis* mesophyll protoplasts of eIF4A1 interaction with the mutated versions of SMG7. Scale bar, 10 μm. **(F)** Fluorescence lifetime measured in a FLIM-FRET interaction assay using the indicated protein combinations transiently expressed in *N. tabacum* leaves (mean, SD, n = 15-26. *\*\*\*\*p* < 0.0001, Student’s t test). **(G)** Alphafold3 model of the SMG7-eIF4A1 interaction interface prediction. SMG7 is represented in grey with the 14-3-3 domain highlighted in green. eIF4A1 is represented in cyan. The right panel shows a close-up view of the interaction. Dashed cyan line represent predicted hydrogen bonds. **(H)** Fluorescence lifetime measured in a FLIM-FRET interaction assay using the indicated protein combinations transiently expressed in *N. tabacum* leaves (mean, SD, n = 20-38. *\*p* < 0.05, *\*\*\*\*p* < 0.0001, Student’s t test).

We next used bimolecular fluorescence complementation (BiFC) in mesophyll protoplasts to dissect the interaction between SMG7 and eIF4A1 and eIF4A2 (hereafter referred to as eIF4A1/2). Co-transfection of nYFP-SMG7 with eIF4A1/2-cYFP produced a strong YFP signal, whereas no signal was detected in the control, where nYFP-SMG7 was co-transfected with another subunit of the eIF4F complex, eIF4B1-cYFP (Figures 1E and S1A). SMG7 consists of an evolutionary conserved N-terminal 14-3-3-like domain and a central helical domain (28), while the C-terminal half is highly diverged and intrinsically disordered (Figure 1D)(22). This region includes a prion-like domain and several motives at the very C-terminus (CD), which are well conserved among plant SMG7 proteins (22, 35). To map the SMG7 region required for interaction with eIF4A1/2, we generated a series of deletion constructs. While deletions spanning the prion-like domain, CD, or the entire C-terminus including both domains produced a BiFC signal, deletion of the 14-3-3 domain entirely abolished the signal (Figures 1E and S1A).

The eIF4A1/2 is uniformly distributed throughout the cytoplasm of mesophyll protoplasts (Figure S1B), whereas the eIF4A1/2-SMG7 BiFC signal appears in distinct cytoplasmic speckles. We have previously shown that SMG7 localizes to P-bodies, and that deletion of its C-terminus (SMG7^Δ702-1059^), which spans the intrinsically disorder regions (IDRs), reduces its partitioning into P-bodies (Figure S1C)(22). This suggests that the majority of SMG7-eIF4A1/2 interaction occurs in P-bodies and reflects the subcellular localization of SMG7. This conclusion is further supported by the observation that the interaction with truncated SMG7^Δ702-1059^ is detectable in speckles as well as in cytoplasm (Figure 1E). We also noticed that the deletion of the N-terminal domain abolished P-body localization of SMG7 (Figure S1C), indicating that its partitioning into P-bodies is primarily mediated by interactions involving the α-helical 14-3-3 and reinforced by the C-terminal IDRs.

The 14-3-3 domain forms a binding pocket that recognizes phosphoserine-residues of phosphorylated UPF1, with the interaction coordinated by evolutionary conserved lysine and arginine residues (27, 28). To determine whether eIF4A1/2 binding to SMG7 occurs through this pocket, we employed Förster resonance energy transfer by fluorescence lifetime microscopy (FLIM-FRET), with SMG7 mutant in the conserved Lys^77^ and Arg^185^ residues (SMG7^K77E^ ^R185E^) known to abolish the phosphoserine binding (27, 28). We observed a shortening of the YFP fluorescence lifetime of SMG7-YFP during co-expression with eIF4A1/2-TagRFP constructs, further confirming their interaction (Figure 1F). In contrast, the fluorescence lifetime remained unchanged when YFP-SMG7^K77E^ ^R185E^ was co-expressed with eIF4A1/2-TagRFP, indicating that these residues are essential for the interaction.

These data indicate that phosphorylation of eIF4A1/2 may facilitate the interaction with SMG7. A search in the PhosPhat 4.0 database (36) for previously identified phosphosites in eIF4A1 and eIF4A2 revealed consistent phosphorylation of Ser^4^ in the N-terminus, and at the Thr^145^ and Ser146 in the N-helicase domain, across several independent datasets. To further explore the interaction interface, we used AlphaFold 3 (AF)(37) to model the SMG7–eIF4A1 complex, incorporating these three phosphorylation sites. The resulting AF model revealed an interaction interface between phosphorylated eIF4A1 and the 14-3-3 domain of SMG7, in which phosphorylated Ser^146^ forms an hydrogen bond with Arg^185^ of SMG7 (Figure 1G), supporting our FLIM-FRET data (Figure 1G). Although Lys^77^ of SMG7 was not predicted to be directly involved in the interaction, its neighbouring residue Tyr^76^, along with the Tyr^186^ forms hydrogen bonds with Ser^146^ of eIF4A1. Thr^145^ of eIF4A1 is not directly part of the interaction interphase in the model, but it may contribute to stabilizing the helicase loop of eIF4A1, which is engaged in the 14-3-3 binding pocket of SMG7 (Figure 1G).

We experimentally validated the AF model using FLIM-FRET and substitutions of Thr^145^ and Ser^146^ in eIF4A1. The interaction with SMG7 was significantly reduced in the non-phosphorylatable eIF4A1^S146A^ mutant and completely abolished when both Thr^145^ and Ser^146^ were mutated (eIF4A1^T145A^ ^S146A^; Figure 1H). In contrast, the phospho-mimetic eIF4A1^S146D^ substitution displayed an interaction strength comparable to that of the wild-type eIF4A1. In conclusion, these results demonstrate that SMG7 interacts with eIF4A1/2, and that this interaction is mediated by the 14-3-3 binding pocket of SMG7 and the Thr^145^-Ser^146^ interface within the N-helical domain of eFI4A. The FLIM-FRET data further suggest that this interaction is facilitated by the phosphorylation at Ser^146^.

### eIF4A1 and eIF4A2 colocalize with SMG7 to P-bodies

We next performed colocalization studies in the roots of *Arabidopsis* lines expressing SMG7-TagRFP (22) along with either eIF4A1-YFP or eIF4A2-YFP. Confocal microscopy showed prominent localization of SMG7-TagRFP to P-bodies (Figure 1A). While the majority of the eIF4A1/2-YFP signal was diffusely distributed throughout the cytoplasm, we also observed distinct foci that colocalized with the larger SMG7-TagRFP marked P-bodies (Figure 2A, Figure S1A). P-bodies are known to increase in size under certain stress conditions, such as heat-shock (23, 38). Indeed, exposing roots to 39°C for 30 min led to a marked increase in both the size and abundance of SMG7-TagRFP labelled P-bodies (Figures 2A, S2A, 2G and S2G). Heat shock also significantly enhanced the partitioning of eIF4A1/2-YFP into cytoplasmic speckles, which co-localized with SMG7-TagRFP. (Figure 2A and S2A).

**Figure 2.**
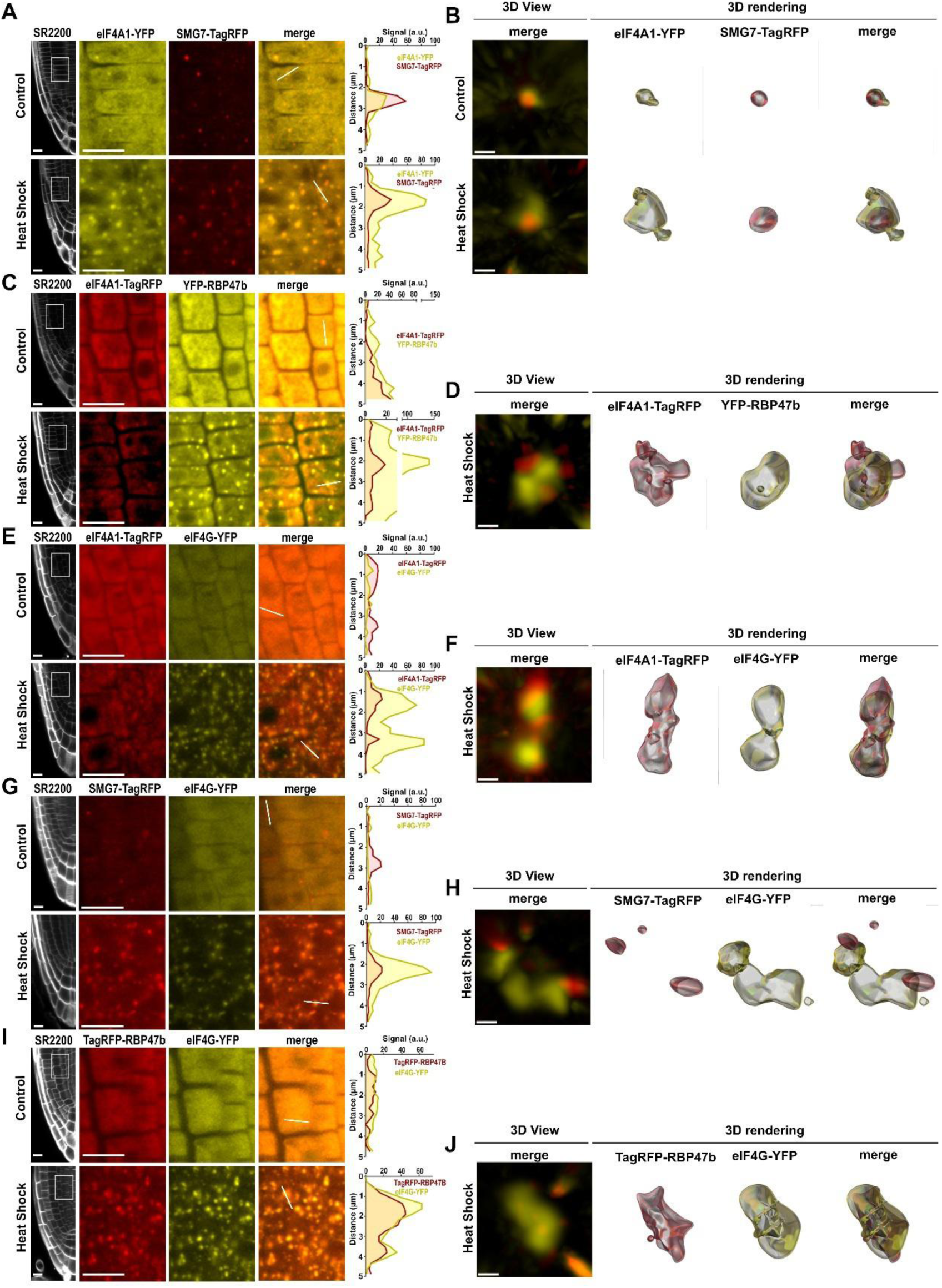
eIF4A1 localizes to P-bodies and SGs in *Arabidopsis* roots (A,C,E,G,I) Confocal micrographs of root cells co-expressing indicated combinations of reporter proteins. Counterstaining with SR2200 dye was used to visualize cell walls. Diagrams on the left show superimposed intensity profiles of YFP and TagRFP signals measured along the lines indicated in the corresponding micrographs. Scale bar = 10 μm. **(B,D,F,H,J)** Super-resolution micrographs of indicated protein condensates visualized by 3D view and 3D rendering using Imaris software. Scale bar = 0.5 μm.

The subunits of the eIF4F typically associate with SGs and *Arabidopsis* eIF4A2 localizes to SGs upon heat shock and salt stress (39). Since SGs frequently associate with P-bodies, we employed structured illumination super-resolution microscopy to distinguish the localization of eIF4A1/2 within these subcellular compartments. Under normal temperature, eIF4A1/2-YFP foci overlapped precisely with the SMG7-TagRFP signal, confirming the localization of eIF4A1/2 to P-bodies. However, following heat shock treatment, the eIF4A1/2-YFP signal occupied a substantially larger volume than SMG7-TagRFP (Figures 2B and S2B), suggesting that in addition to localizing to P-bodies, heat shock promotes expansion of eIF4A1/2 condensation beyond P-bodies, likely into associated SGs.

To test this possibility, we generated double reporter lines expressing YFP-RBP47b, an *Arabidopsis* SG marker (23, 38), and eIF4A1-TagRFP or eIF4A2-TagRFP. Under normal conditions, YFP-RBP47b exhibits diffused cytoplasmic localization, but partitions into prominent SGs upon heat shock (Figure 2C). Super-resolution microscopy revealed that the heat-induced YFP-RBP47b signal partially overlaps with the eIF4A1/2-TagRFP (Figures 2D and S2D). Importantly, co-localization analysis of SGs and PBs in heat-stressed plants expressing YFP-RBP47b and SMG7-TagRFP showed that while these two proteins form closely associated granules, their signals do not overlap and occupy distinct territories (Figure S2E and S2F). These data indicate that under heat shock conditions, eIF4A1/2 localize to both SGs and P-bodies.

We next sought to determine whether localization to P-bodies is specific to eIFA1/2, or whether it represents a general feature of the eIF4F complex in plants. Similar to YFP-RBP47b, the eIF4G-YFP reporter (22) displayed diffuse cytoplasmic localization in root cells under normal conditions, but re-localized to cytoplasmic speckles following heat-shock treatment (Figures 2E,G,I). Interestingly, despite being part of the same complex, eIF4G speckles only partially overlapped with those of eIF4A1/2 (Figures 2E and S2H). While heat-induced eIF4G speckles associated with SMG7-marked P-bodies, they formed distinct subdomains (Figures 2G, 2H), suggesting that eIF4G partitions into SGs. This was confirmed in the double reporter line expressing TagRFP-RBP47b and eIF4G-YFP, where both makers co-localized in the same heat-induced granules (Figure 2J and 2I).

In conclusion, these co-localization experiments show that eIF4A1/2 have the capacity to partition into both P-bodies and SGs, a feature that distinguishes them from their interaction partner eIF4G, which localizes exclusively to SGs under stress conditions.

### eIF4A1/2 limit the condensation of SGs and P-bodies in *Arabidopsis* root cells

In addition to its role in translation, eIF4A functions as an RNA chaperone reducing SG condensation in human cells by limiting RNA-RNA interactions (6). To determine whether eIF4A1/2 influences SG dynamics in plants, we analysed the condensation of RBP47b in *eif4a1* and *eif4a2* mutants following heat shock treatment. Both mutations resulted in an increase in the size of the RBP47b-labeled condensates, as well as a greater total condensate signal per unit of cell volume (Figures 3A-3C). These results indicate that eIF4A1/2 limit SG condensation during heat stress in *Arabidopsis*.

**Figure 3.**
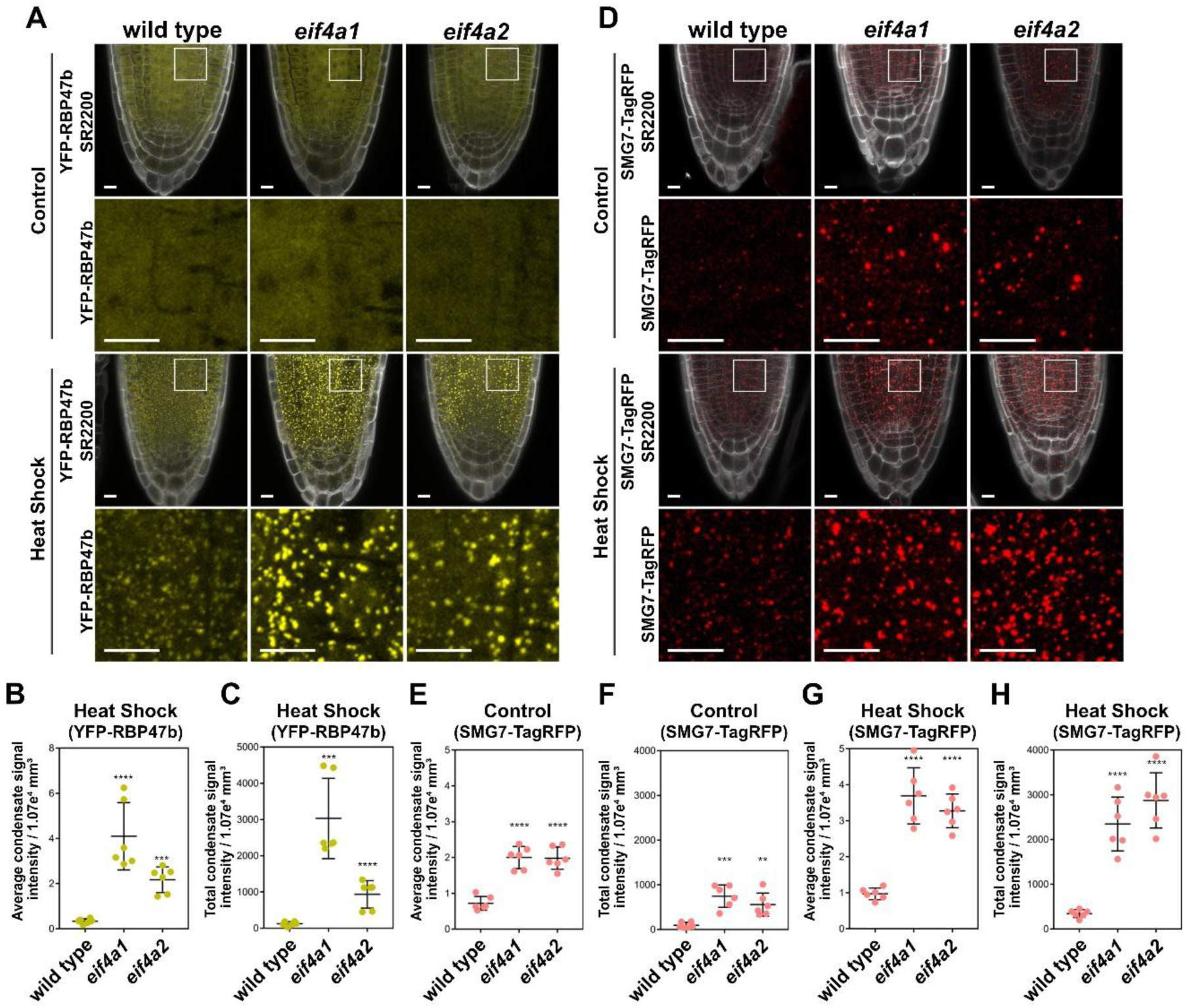
eIF4A1/2 limit the condensation of SGs and P-bodies in *Arabidopsis* roots. **(A)** Confocal micrographs of *Arabidopsis* roots from wild type, and *eif4a1* and *eif4a2* mutants expressing YFP-RBP47b, under heat shock and control conditions. Cell walls were visualized with SR2200 counterstaining. Scale bars = 10 μm. **(B)** Dot plots showing average signal intensity per condensate and **(C)** total condensate signal intensity of all SGs detected within the volume of 1.07e^4^ μm^3^ encompassing endodermic and cortical cells (n = 6; \*\**\*p* < 0.001, *\*\*\*\*p* < 0.0001, Student’s t test). **(D)** Confocal micrographs of *Arabidopsis* roots expressing SMG7-TagRFP, under control conditions and heat shock. Cell walls were visualized with SR2200 counterstaining. Scale bars = 10 μm. **(E,G)** Dot plots showing average signal intensity per condensate and **(F,H)** total condensate signal intensity of all P-bodies detected within the volume of 1.07e^4^ µm^3^ encompassing endodermic and cortical cells. (n = 6; \*\**\*p* < 0.001, *\*\*\*\*p* < 0.0001, Student’s t test).

We next investigated whether eIF4A1/2 also modulate P-body condensation by examining SMG7-TagRFP signal in the *eif4a1* and *eif4a2* mutants (Figure 2D-2H). Under standard conditions, both the average size of P-bodies and their total signal intensity per cell volume were elevated in the mutants (Figures 2D-2F). This effect was even more pronounced following heat shock, where deficiency in either eIF4A isoform led to a marked increase in P-body signal compared to wild type (Figures 2D,2G and 2H). Together, these findings suggest that the RNA helicases eIF4A1 and eIF4A2 act to restrain the condensation of both P-bodies and SGs in *Arabidopsis*, supporting a conserved RNA chaperone function across eukaryotes.

### eIF4A1 and eIF4A2 localize to meiotic P-bodies and modulate their size

We previously demonstrated that efficient partitioning of SMG7 into P-bodies is essential for termination of meiosis and the transition to post-meiotic development of *Arabidopsis* microspores (22). Given the influence of eIF4A1/2 on P-body size and their physical interaction with SMG7, we investigated their function during male meiosis. Confocal microscopy of anthers from *eIF41:TagRFP* and *eIF4A2:TagRFP* plants revealed strong signal in the tapetum, the innermost cell layer of the anther locule, likely reflecting a high rate of translation in these metabolically highly active cells (Figures 4A and S3A). In meiocytes, eIF4A1/2 formed distinct foci that were most prominent during pachytene (Figures 4A and S3A) and co-localized with SMG7 (Figured 4B, 4C, S3B, and S3C). Notably, the SMG7-marked P-bodies in pachytene-stage meiocytes were significantly larger than those observed in root and tapetal cells (Figure S3).

**Figure 4.**
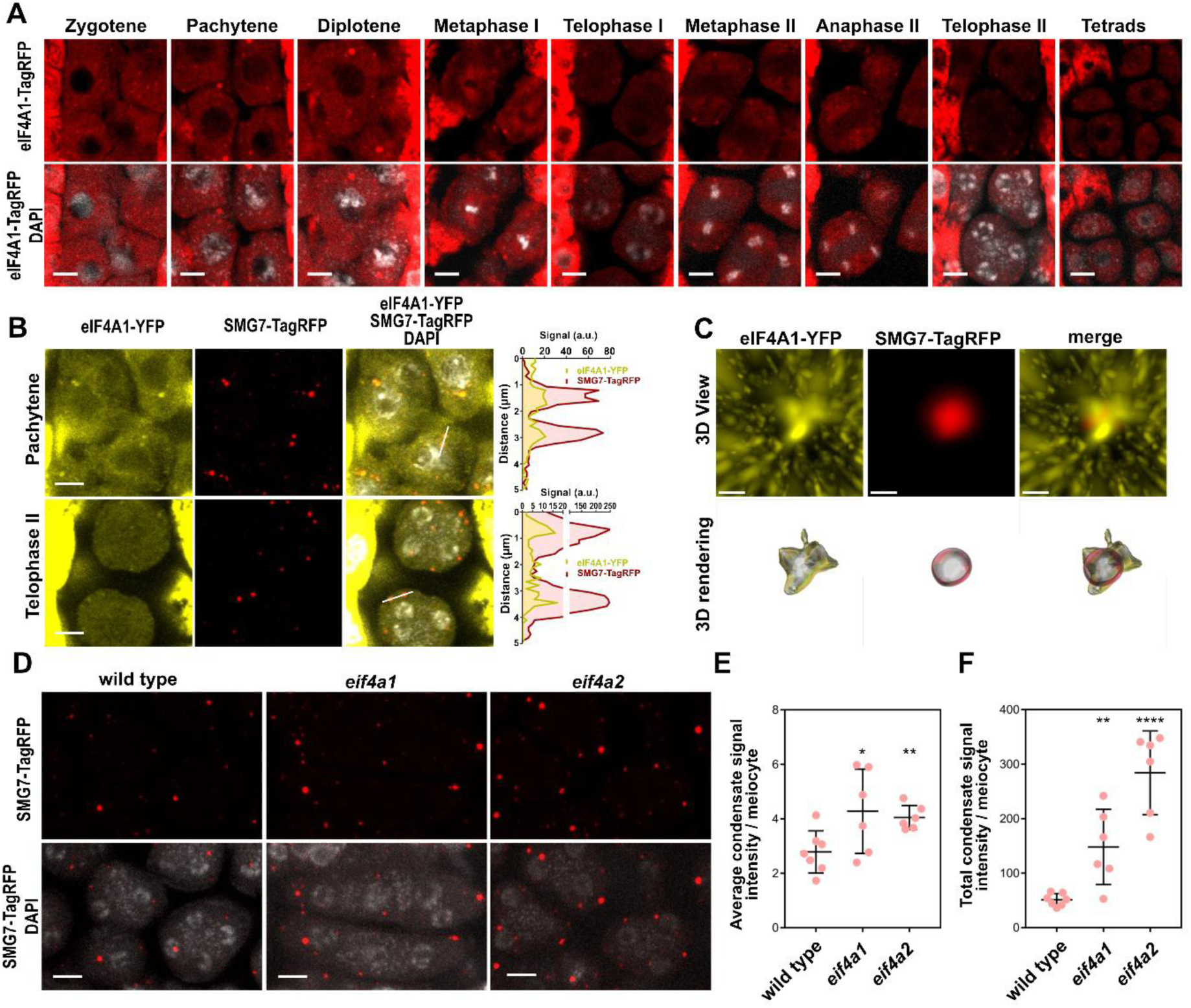
eIF4A1/2 localize to meiotic P-bodies and modulate their condensation. **(A)** Confocal micrographs of *Arabidopsis* anthers showing localization of eIF4A1-TagRFP in the course of meiosis. DNA is counterstained with DAPI. Scale bars = 5 μm. **(B)** Confocal micrographs of *Arabidopsis* meiocytes co-expressing eIF4A1-TagRFP and SMG7-TagRFP. Diagrams on the left show superimposed intensity profiles of YFP and TagRFP signals measured along the lines indicated in the corresponding micrographs. Scale bar = 10 μm. **(C)** Super-resolution micrographs of indicated protein condensates visualized by 3D view and 3D rendering using Imaris software. Scale bar = 0.5 μm. **(D)** Confocal micrographs of *Arabidopsis* meiocytes of wild type, *eif4a1* and *eif4a2* plants expressing SMG7-TagRFP. Scale bars, 5 μm. **(E)** Dot plots showing average signal intensity per condensate in a meiocyte and **(F)** total condensate signal intensity in a meiocyte (n = 6 meiocytes; \*\**\*p* < 0.001, *\*\*\*\*p* < 0.0001, Student’s t test).

The strong partitioning of eIF4A1/2 to meiotic P-bodies suggests a regulatory role in maintaining their size homeostasis. To test this, we analysed the condensation levels of SMG7-marked P-bodies in meiocytes of wild-type, *eif4a1*, and *eif4a2* mutants. Consistent with observations in root cells, P-bodies in meiocytes lacking either helicase were significantly larger and more abundant compared to those in the wild type (Figures 4D-4F).

### Meiotic bodies

Meiotic P-bodies exhibit distinct characteristics compared to those observed in somatic cells. Notably, their larger size (Figures S3D-S3F) and enhanced recruitment of eIF4A1/2 (Figures 4A and S3A) resemble the P-bodies formed in response to heat shock in root cells. This prompted us to investigate whether these meiotic condensates also associate with SG markers. Strikingly, confocal microscopy of anthers from the *YFP:RBP47b* line revealed prominent cytoplasmic speckles in meiocytes at normal temperature, whereas the YFP-RBP47b signal appeared more diffuse in the surrounding tapetal cells (Figure 5A). Nucleation of YFP-RBP47b speckles was observed throughout all meiotic stages, beginning in late leptotene, becoming particularly prominent during zygotene and pachytene, and remaining visible through the tetrads stage (Figure 5A). Although heat shock increased the number of RBP47b speckles in meiocytes, their size and signal intensity remained comparable to those observed at control temperature (Figures 5B-5E).

**Figure 5.**
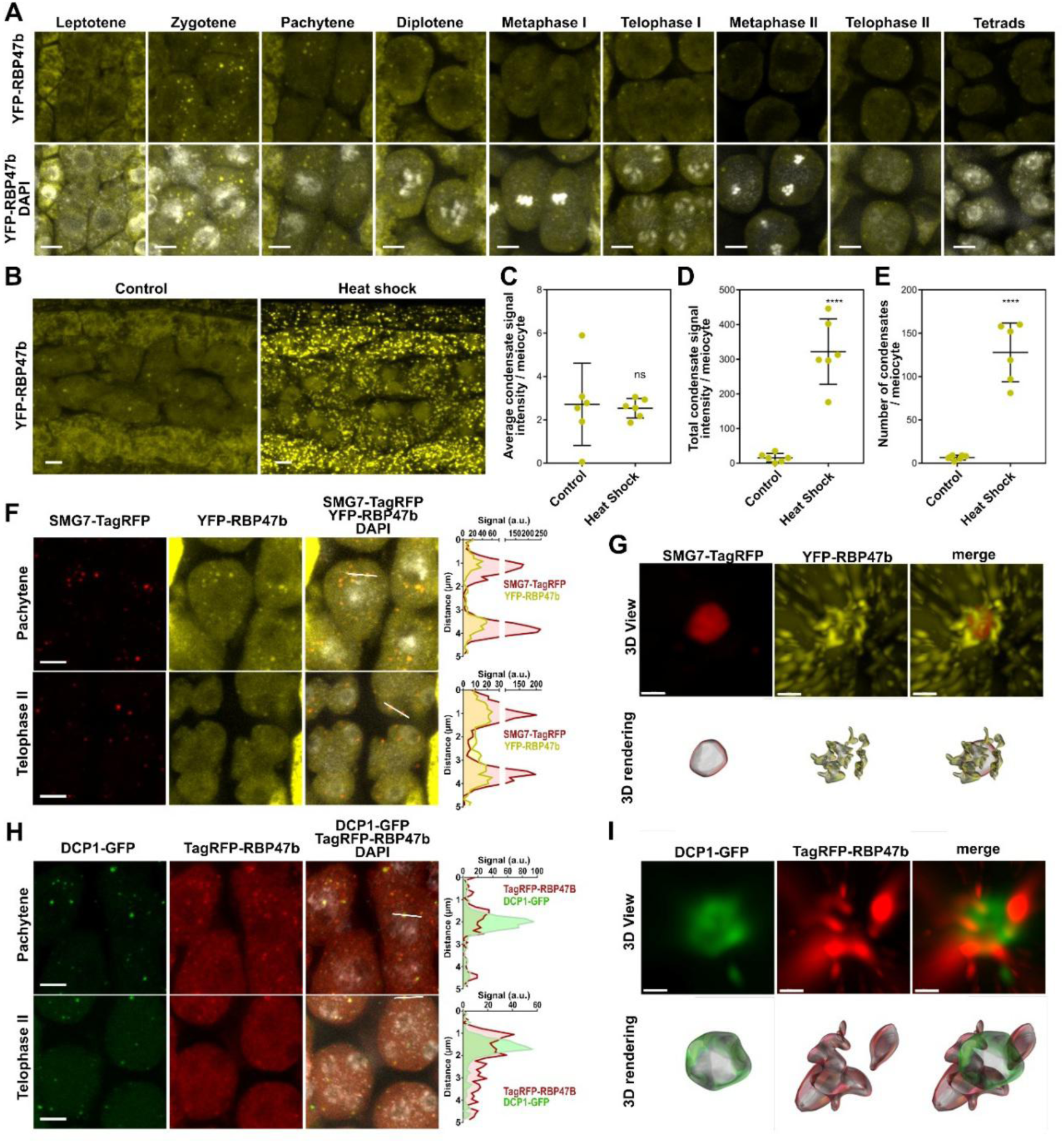
Composition of M-bodies. **(A)** Confocal micrographs of *Arabidopsis* pollen mother cells depicting the expression and localization of YFP-RBP47b in the course of meiosis. DNA is counterstained with DAPI. Scale bars = 5 μm. **(B)** Confocal micrographs of anther lobes, showing expression and localization of YFP-RBP47b in pachytene meiocytes and the surrounding tapetum upon heat shock or under control conditions without heat shock application. Scale bar = 5 μm. **(C-E)** Dot plots showing average of the signal intensity in condensates from a meiotic cells (C), total signal intensity in all condensates per meiotic cell (D), and the average number of condensates per meiocyte (E), (n = 6; *ns* ≥ 0.05, *\*\*\*\*p* < 0.0001, Student’s t test). **(F)** Confocal micrographs of *Arabidopsis* meiocytes co-expressing SMG7-TagRFP and YFP-RBP47b. Diagrams on the left show superimposed intensity profiles of YFP and TagRFP signals measured along the lines indicated in the corresponding micrographs. Scale bar = 5 μm. **(G)** Super-resolution micrographs of SMG7-TagRFP / YFP-RBP47b condensates visualized by 3D view and 3D rendering using Imaris software. Scale bar = 0.5 μm. **(H)** Confocal micrographs of *Arabidopsis* meiocytes co-expressing DCP1-GFP and YFP-RBP47b. Diagrams on the left show superimposed intensity profiles of GFP and TagRFP signals measured along the lines indicated in the corresponding micrographs. Scale bar = 5 μm. **(G)** Super-resolution micrographs of DCP1-GFP / YFP-RBP47b condensates visualized by 3D view and 3D rendering using Imaris software. Scale bar = 0.5 μm.

While confocal microscopy showed co-localization of RBP47b and SMG7 speckles (Figure 5F), super-resolution imaging revealed a more nuanced organization: RBP47b forms a shell around an SMG7-enriched core, with the two regions occupying distinct, non-overlapping domains (Figure 5G). In these lines, RBP47b is expressed in meiotic cells under the control of the strong constitutive *RPS5a* promoter. To rule out the possibility that RBP47b condensation in meiocytes is an artifact of overexpression, we validated this observation using two additional SG marker lines, eIF4G-YFP and UBP1b-YFP, each expressed from their native promoters (Figure S4). Both SG components formed distinct cytoplasmic foci in meiocytes under standard growth conditions, and their abundance markedly increased upon heat shock. These foci co-localized with SMG7 bodies, but as seen with RBP47b, they formed a peripheral shell around the SMG7 core without signal overlap (Figures S4C-S4F).

To further confirm these observations, we examined the spatial relationship between RBP47b and DCP1, a core component of P-bodies. In *Arabidopsis* lines co-expressing TagRFP-RBP47b and DCP1-YFP, RBP47b was again found to condense at the surface of DCP1-labeled speckles, forming a distinct outer layer (Figures 5H and 5I). In conclusion, these localization experiments demonstrate that *Arabidopsis* meiocytes contain prominent cytoplasmic speckles composed of a P-body core surrounded by a shell of SG components. We propose the term Meiotic bodies (M-bodies) to describe this specialized class of RNP granules.

### SMG7 recruits eIF4A1/2 into the core of M-bodies

The eIF41/2 helicases localize to both P-bodies and SGs in somatic cells under heat shock (Figures 2D, 2F, S2D and S2H). To further refine their location within M-bodies, we performed colocalization analysis of eIF4A1/2 with DCP1 and the SG markers RBP47b, eIF4G and UBP1b. Similar to SMG7, the eIF4A1/2-TagRFP signal overlapped with DCP1 (Figure 6A,6B, S6A and S6B) and extended only slightly beyond the DCP1-defined volume (Figures 6B and S6B). In contrast, super-resolution microscopy showed that eIF4A1/2 occupied adjacent, but not overlapping territories relative to RBP47b, eIF4G and UBP1, which define the M-body shell (Figures 6C, 6D, and S5C-H). This finding was unexpected, given that eIF4A1/2 localize to SGs during heat shock and physically interact with eIF4G as part of the eIF4F complex. These data indicate that in meiotic cells eIF41/2 helicases are restricted, along with other P-body components, to the core of M-bodies.

**Figure 6.**
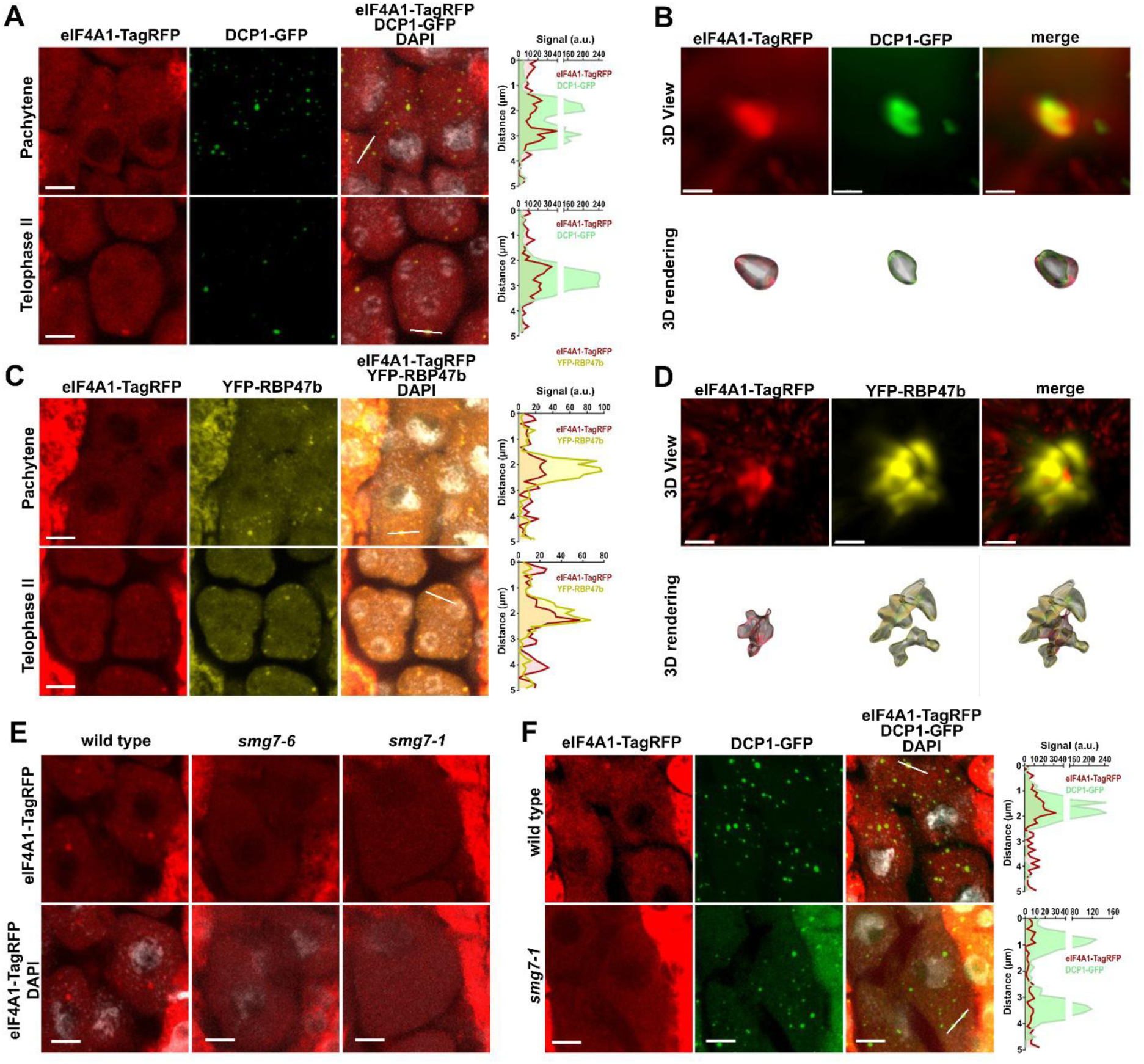
Localization of eIF4A1 in M-bodies. **(A)** Confocal micrographs of *Arabidopsis* meiocytes co-expressing eIF4A1-TagRFP and DCP1-GFP. Diagrams on the left show superimposed intensity profiles of GFP and TagRFP signals measured along the lines indicated in the corresponding micrographs. Scale bar = 5 μm. **(B)** Super-resolution micrographs of eIF4A1-TagRFP / DCP1-GFP condensates visualized by 3D view and 3D rendering using Imaris software. Scale bar = 0.5 μm. **(C)** Confocal micrographs of *Arabidopsis* meiocytes co-expressing eIF4A1-TagRFP and YFP-RBP47b. Diagrams on the left show superimposed intensity profiles of YFP and TagRFP signals measured along the lines indicated in the corresponding micrographs. Scale bar = 5 μm. **(D)** Super-resolution micrographs of eIF4A1-TagRFP / YFP-RBP47b condensates visualized by 3D view and 3D rendering using Imaris software. Scale bar = 0.5 μm. **(E)** Confocal micrographs of *Arabidopsis* meiocytes of wild type, *smg7-6* and *smg7-1* plants expressing eIF4A1-TagRFP. Scale bars, 5 μm. **(F)** Confocal micrographs of *Arabidopsis* meiocytes of wild type and *smg7-1* mutants co-expressing eIF4A1-TagRFP and DCP1-GFP. Scale bars, 5 μm. DNA in (A), (C), (E) and (F) is counterstained with DAPI.

Several members of the DEAD-box helicase family associate with RNP granules and contain IDRs that facilitate their phase separation into condensates (12, 40, 41). However, eIF4A1/2 lack IDRs and their yeast orthologue has been shown to fail in forming phase-separated droplets *in vitro* (12), raising the question of how they are recruited into M-bodies. SMG7 has previously been reported to mediate the translocation of UPF1 and TDM1 into P-bodies (22, 26). Therefore, we next investigated whether SMG7 is required for eIF4A1/2 M-body localization. To this end, we crossed *eIF4A1-TagRFP* and *eFF4A2-TagRFP* reporters to *smg7-1* null mutants as well as to *smg7-6* hypomorphic allele, which encodes a truncated SMG7 protein lacking the C-terminal IDR (Figure 1D) and exhibits reduced partitioning into P-bodies (Figure S1C) (22). The loading of eiF4A1/2 in cytoplasmic speckles was significantly reduced in *smg7-6* and completely abolished in *smg7-1* mutants (Figures 6E and S6I).

To exclude the possibility that M-bodies fail to nucleate in the absence of SMG7, we analysed the co-localization of eIF4A1 with DCP1 in *smg7-1* mutants. This analysis confirmed that M-bodies do nucleate without SMG7, but they do not incorporate eIF4A1. Together with our interaction data (Figure 1), these findings indicate that SMG7, via its 14-3-3 domain, binds eIF4A1/2 and is responsible for recruiting them to the core of M-bodies.

### Downregulation of eIF4A1 enhances TDM1 localization to M-bodies and facilitates meiotic exit

We have previously shown that M-bodies play a crucial role in the termination of meiosis and the cell-fate transition to post-meiotic microspore development. Central to this transition is the SMG7-mediated recruitment of TDM1 into P-bodies during meiosis II (22). In telophase II, TDM1 co-localizes with SMG7, RBP47b and eIF4A1 as observed by confocal microscopy, further confirming its presence in the M-body (Figure S7). To more precisely determine the localization of TDM1 within the M-body, we performed super-resolution microscopy. This analysis revealed that TDM1-YFP occupies the same volume as SMG7 and eIF4A1/2 in the M-body core and, like eIF4A1/2, extends slightly beyond the SMG7-defined region (Figure 7A and 7B). Consistent with this, the TagRFP-RBP47b signal marking the M-body shell was found to surround the TDM1-YFP signal (Figure 7C).

**Figure 7.**
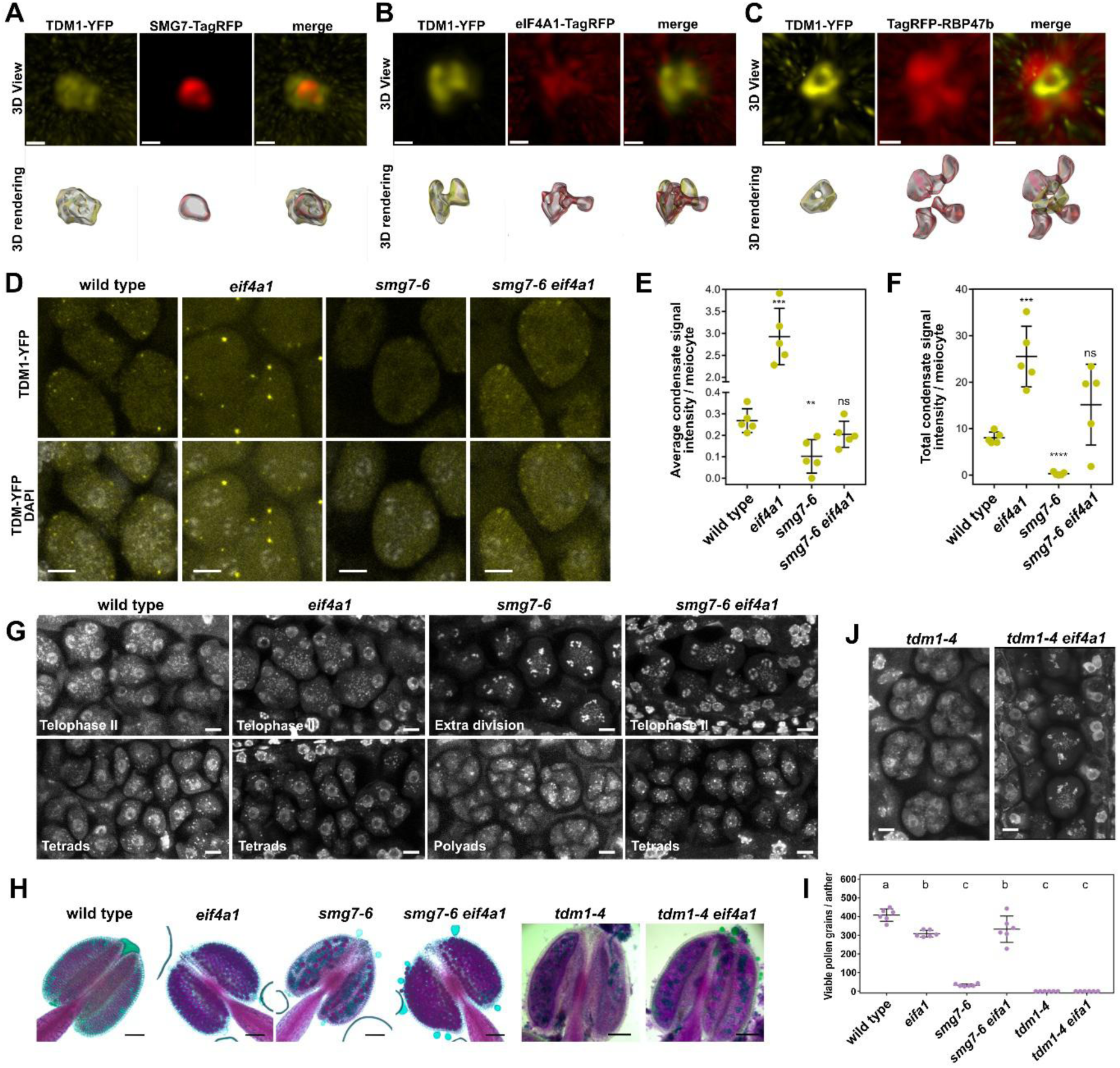
Effect of eIF4A1 on TDM1 (A -C) Super-resolution micrographs of indicated protein condensates visualized by 3D view and 3D rendering using Imaris software. Scale bar = 0.5 μm. **(D)** Confocal micrographs of pollen mother cells in telophase II of wild type, *eif4a1*, *smg7-6* and *smg7-6 eif4a1* plants expressing TDM1-YFP. Scale bars = 5 μm. **(E)** Dot plots indicating average of the signal intensity of condensates and **(F)** total signal intensity of all condensates in meiotic cells from (D); (n = 5 meiocytes; *ns* ≥ 0.05, *\*\*p* < 0.01, *\*\*\*p* < 0.001, Student’s t test). **(G)** Confocal micrographs of meiocytes of wild type, *eif4a1*, *smg7-6* and *smg7-6 eif4a1* plants, stained with DAPI. Scale bars = 5 μm. **(H)** Alexander staining of anthers for pollen viability of wild type and indicated mutants. Scale bars = 100 μm. **(I)** Dot plots showing quantification of the viable pollen grains wild type and indicated mutants (n = 6. One-way ANOVA followed by Tukey’s post-hoc test *p* < 0.05). (J) Confocal micrographs of meiocytes of *tdm1-4* and *tdm1-4 eif4a1* plants stained with DAPI. Scale bars = 5 μm.

Our data indicate that eIF4A1/2 influences the size of M-body, as assessed by SMG7 signal intensity (Figures 4D-4F). Since SMG7 recruits TDM1 into M-bodies, we next asked whether eIF41/2 also affects the partitioning of TDM1 into these structures. To test this, we introduced the TDM1-YFP marker into the *eif4a1* mutant background and analysed TDM1 localization in M-bodies. Strikingly, the absence of eIF4A1 led to a substantial increase in TDM1 signal within M-bodies (Figures 7D-7F), an effect that appears even more pronounced than the SMG7 signal increase in the same mutant background (Figure 4D-4F). The TDM1-recruitment into M-bodes is severely impaired in *smg7-6* mutants (Figure 7D-7F) (22) due to inefficient M-body partitioning of the C-terminally truncated SMG7 protein (Figure S1C). Importantly, TDM1 localization to M-bodies is partially restored in *smg7-6 eif4a1* double mutants (Figures 7D-F), suggesting that eIF4A1 antagonizes the recruitment or retention of TDM1 in M-bodies.

*Arabidopsis smg7-6* mutants fail to terminate meiosis due to inefficient recruitment of TDM1 in M-bodies (22). Upon formation of haploid nuclei in telophase II, *smg7-6* meiocytes attempt additional rounds of chromosome segregations, leading to unequal chromosome distribution, the formation of polyads instead of tetrads, and a reduction of viable microspores and pollen (22, 42). Since the eIF4A1-deficiency restores TDM1 recruitment to M-bodies in *smg7-6* mutants (Figure 7D-F), we investigated whether this also restores normal meiotic progression and pollen formation. Indeed, while *smg7-6* mutants display extra meiotic divisions, polyads formation, and a reduced set of pollen, *smg7-6 eif4a1* double mutants form regular tetrads consisting of four haploid nuclei and exhibit restored pollen counts (Figure 7G-I).

TDM1 localized to M-bodies promotes meiotic exit by sequestering eIFiso4G2 and inhibiting translation. As eIF4A1 is a translation initiation factor, it is possible that the *eif4a1* mutation rescues the *smg7-6* phenotype directly by reducing translation, independently of TDM1 localization. To test this, we analyzed meiosis and fertility in *eif4a1 tdm1* double mutants. Like *smg7-6*, *tdm1* mutants exhibit extra meiotic divisions and form polyads, leading to pollen abortion (22, 43). However, this phenotype was not rescued in *eif4a1 tdm1* double mutants, which remained infertile (Figures 7H-7J). These results suggest that eIF4A1 deficiency restores fertility in *smg7-6* mutants by enhancing TDM1 retention in M-bodies, rather than through a general impairment of translation.

## Discussion

SMG7 is a prominent component of P-bodies, but the functional significance of this localization remains unclear. SMG7 is primarily known for its role in NMD where it promotes RNA degradation following target recognition in both animals and plants (44–46). It binds to phosphorylated UPF1 through its 14-3-3, but the molecular events that occur downstream of this interaction are less well understood. The ability of SMG7 to translocate UPF1 into P-bodies initially led to the hypothesis that P-bodies serve as sites of RNA degradation (26, 27, 47). However, subsequent studies demonstrating that NMD can occur independently of P-bodies challenged this model (48, 49).

In our previous work, we described an NMD-independent function of SMG7 in meiotic exit. This mechanism relies on the interaction between SMG7 and TDM1, which similarly to UPF1 occurs through the 14-3-3 phosphoserine-binding pocket, and the ability of SMG7 to translocate TDM1 into the P-body core of M-bodies (22)(Figure 7). Importantly, *smg7-6* mutants, in which SMG7 retains the ability to bind TDM1 but has reduced capacity to localize into P-bodies, exhibit a phenotype similar to *tdm1* null mutants. These findings indicate that the key function of SMG7 in meiotic exit is to transport or sequester TDM1 into M-bodies. Here we identified eIF4A1/2 as additional SMG7 interactors that bind via its 14-3-3 phosphoserine-binding pocket (Figure 1). Our data further show that SMG7 is necessary for the localization of eIF4A1/2 to the P-body core of M-bodies (Figure 6). As with TDM1, C-terminal truncation of SMG7 does not disrupt the interaction with eIF4A1/2, but substantially reduces their recruitment to M-bodies, highlighting the importance of the disordered C-terminal region for P-body localization. SMG7 partitioning into P-bodies also requires its structured 14-3-3 domain (Figure S1), but it is independent of the phosphoserine-binding pocket (22) suggesting that the 14-3-3 domain contributes to both substrate binding and P-body association through distinct interactions. A similar dual role for the 14-3-3 domain in substrate binding and biocondensate formation has also been reported for the 14-3-3ζ protein (50). Together, these observations define a novel role for SMG7 as an adaptor or scaffold protein that facilitates the recruitment or retention of specific client proteins in P-bodies.

The eIF4A1/2 helicases are core factors of the translation initiation complex eIF4F. Recently, eIF4A was proposed to act independently of its function in translation initiation as an RNA chaperone that limits condensation of SGs in human cells (6). We found that inactivation of either of the *Arabidopsis* eIF4A isoform leads to increased size of SGs in root cells (Figure 3), suggesting a conserved role of eIF4A in SG homeostasis. Unlike in human cells, where eIF4A is present in excess over the other eIF4F subunits (6), quantitative proteomics across different *Arabidopsis* tissues show that eIF4A1/2 are roughly equimolar to eIF4G and its isoforms eIFiso4G1/2 (51). Thus, despite genetic redundancy (33), loss of a single eIF4A orthologue can cause insufficiency and profound effect on SG formation.

We also observed that eIF4A1/2 are constitutively present in P-bodies, a finding not previously reported. Although the eIF4E cap-binding protein associates with both SGs and P-bodies, other eIF4F components are typically restricted to SGs (52, 53). Several lines of evidence suggest that eIF4A1/2 regulate P-body dynamics as an RNA chaperone. First, eIF4A1/2 localization does not coincide with that of eIF4G (Figures 2 and S2). While both co-localize in SGs during heat stress, eIF4G remains confined to SGs, whereas eIF4A1/2 are also present in adjacent P-bodies. This distinct compartmentalization is even more pronounced in M-bodes, where eIF4G localizes to the shell, while eIF4A1/2 are restricted to the P-body-like core (Figure S5), indicating an eIF4F independent function of eIF4A1/2. Second, *eif4a* mutants show enlarged P-bodies in roots and increased size of the P-body core in M-bodies, consistent with a role in counteracting the RNP condensation. Finally, *eif4a1* mutants exhibit enhanced partitioning of TDM1 into M-bodies, which can rescue meiotic exit in the hypomorphic *smg7-6* background (Figure 7). This could reflect stronger binding of TDM1 to a P-body constituent such as SMG7, or increased retention of RNPs subassemblies within P-bodies. The presence of larger M-bodies in *eif4a* mutants (Figure 5) supports the latter, although decreased competition with eIF4A1/2 for SMG7 binding may also contribute to increased TDM1 recruitment.

Given that SMG7 recruits eIF4A1/2 to the P-body core of M-bodies, and that eIF4A1/2, in turn, limit P-body condensation and thereby restrict further partitioning of SMG7, the SMG7-eIF4A interaction module may constitute a negative feedback loop that maintains P-body size homeostasis. In this model, an increase in P-body size would lead to greater SMG7 accumulation, which recruits more eIF4A. The resulting increase in eIF4A activity would counteract further condensation, thereby limiting P-body growth. Our data indicate that the SMG7-eIF4A interaction is influenced by eIF4A phosphorylation (Figure 1), and plant eIF4As have been reported to undergo phosphorylation in response hypoxia and heat stress (54, 55). Thus, the SMG7-eIF4A module may dynamically adjust the extend of P-body condensation in response to cellular conditions, thereby affecting RNA processing and gene expression under varying physiological stimuli. Other P-body associated DEAD-box helicases from the DDX family likely contribute to this process as they have been implicated in P-body assembly and RNA turnover (8, 12, 40).

Indeed, while P-bodies are generally a constitutive feature of plant cells, they are dynamic structures that vary in size and composition depending on cellular and environmental context (23, 56, 57). Here we show that pollen mother cells contain a distinct type of RNP granule, termed M-bodies, consisting of a P-body core that is larger than P-bodes in somatic cells, and a shell enriched in SG-specific proteins (Figure S7D). Partitioning of eIF4A1/2 into M-bodies is also more pronounced than their localization to P-bodies in root cells. This may be partly due to regulation of eIF4A by cyclin-dependent kinase 1 (CDKA;1), which phosphorylates eIF4A1 and renders it translationally inactive (58). Since CDKA;1 is the key driver of meiotic progression in *Arabidopsis* (59, 60), its activity may increase the pool of translationally inactive eIF4A1 available for SMG7 interaction and recruitment to M-bodies.

In our previous work we showed that M-bodies play an important role in meiotic exit. They change their composition by recruiting TDM1 during meiosis II, which interacts and temporarily sequesters eIFisoG2. This is thought to halt translation and promote meiotic exit (22). Here we show the eIF4A1/2 contributes to this M-body remodelling by regulating TDM1 recruitment (Figure 7). Notably, the colocalization of SG and P-body markers has also been observed in rice, suggesting that M-bodies represent a conserved feature of plant meiosis (61). In rice, these condensates sequester MEL2, an RNA binding protein required for early meiotic progression, further underscoring importance of M-bodies in regulating meiotic events across plant species.

M-bodies may also function in the storage and translational repression of mRNAs transcribed early in meiosis, but destined for expression only during later stages or post-meiotically. Similar mechanisms have been described in yeast and human oocytes, where late meiotic and post-meiotic transcripts are sequestered in cytoplasmic RNP condensates nucleated by dedicated RNA-binding proteins (62, 63). Single-cell transcriptomic studies in maize suggest that major transcriptional changes occur during early meiosis, and some of the transcripts persist through to post-meiotic pollen development (64, 65), implying a need for their translational repression and storage. The structure of M-bodies is reminiscent of the of P-body and SG associations formed during heat shock (Figure 2), and may similarly reflect a high concentration of translationally repressed mRNA in pollen mother cells. In this context, the partitioning of eIF4A into M-bodies could influence the dynamics and translatability of stored mRNAs, similar to the role described for the DD3X helicase in SGs (11).

The discovery of M-bodies as distinct RNP granules with both P-body and SG-like features highlights the complexity of RNA regulation in reproductive cells, suggesting that post-transcriptional control via biomolecular condensates plays a central role in plant meiosis. These insights open new avenues for investigating how cells spatially and temporally coordinate gene expression through dynamic condensate architecture.

## Materials and Methods

### Plant material and growth conditions

*Arabidopsis thaliana* (ecotype Col-0), mutant lines and transgenic lines were grown on soil in growth chambers at 21°C at 50-60% of humidity under 16/8 h light/dark cycles. Roots were analysed from 4 days old seedlings grown in agar plats (0.8% w/v, pH 5.8) supplemented with half-strength Murashige and Skoog medium (½ MS). Heat shock was applied to roots by immersing sealed plates in a water bath tempered at 39°C for 30 min and rapidly processed for fixation. Heat shock was applied to meiotic anthers by immersing whole inflorescence in liquid ½ MS medium supplemented with 5% sucrose in 1.5 ml tube, and incubation in a heat block at 39°C for 20 min.

The following *Arabidopsis* mutant lines were used in this study: *smg7-1* (29), *smg7-6* (30), *tdm1-4* (42), eif4a1 and eif4a2 (33). The mutant line *ubp1b-1* was obtained from NASC (Gabi-Kat GK-262E01-014951) and the primers UBP1b.BamHI.V2.F and UBP1b.6.R (Table S2) were used for PCR-genotyping. The following reporter *Arabidopsis* lines were used: SMG7-MYC (30, 32), eIF4G-YFP (22), SMG7-TagRFP (22), and TDM-YFP (22). These reporter lines were generated in this study: *eIF4A1-YFP*, *eIF4A2-YFP*, *eIF4A1-TagRFP*, *eIF4A2-TagRFP*, *YFP-RBP47b*, *TagRFP-RBP47b*, *UBP1b-YFP*, *UBP1b-TagRFP* and *DCP1-GFP*.

### Plasmid construction and generation of transgenic *Arabidopsis* lines

Primer sequences for plasmid construction are listed in Table S2. The constructs described in the following section were generated using the Gateway® cloning technology (Invitrogen) and the destination vectors from the pGWB series (66). The genomic regions containing the putative promoters and ORFs without stop codons of *eIF4A1*, *eIF4A2* and *DCP1* were amplified with the primers eIF4A1.Prom.F and eif4A1.nostop.r, eif4A2.Promoter.Topo.F and eiF4A2.nostop.r, DCP1_FP and DCP_RP. The fragments were introduced into pENTR™/D-TOPO using blunt-end TOPO® Cloning reactions, and after LR recombination into pGWB640, pGWB659 or pGWB650, the final constructs pGWB640-eIF4A1, pGWB659-eIF4A1, pGWB640-eIF4A2, pGWB659-eIF4A2, and pGWB650-DCP1 were generated to create the lines *eIF4A1-YFP*, *eIF4A2-YFP*, *eIF4A1-TagRFP*, *eIF4A2-TagRFP* and *DCP1-G3GFP*, respectively. These lines were generated transforming the corresponding mutant lines and using *Agrobacterium tumefaciens* and the floral dip method.

The genomic region containing the putative promoter and ORF without stop codon of *UBP1b* was amplified with the primers UBPb1.EcoRI.F.V1 and UBPb1EcoRV.R.V2. The fragment was cloned into the *EcoR*I and *EcoR*V sites of pENTR™11, and LR-recombined into pGWB640 and pGWB659 to obtain the constructs pGWB640-UBP1b and pGWB659-UBP1b, respectively. The constructs pGWB640-UBP1b and pGWB659-UBP1b were used to generate the lines *UBP1b-YFP* and *UBP1b-TagRFP*, respectively, by transforming the mutant line *ubp1b-1* and using *A. tumefaciens* and the floral dip method.

The genomic region containing the ATG and the stop codons of RBP47b was amplified with the primers RBP47b_cacc_topo_F_V2 and RBP47b_stop_R. The fragment was introduced into pENTR™/D-TOPO using blunt-end TOPO® Cloning reaction, obtaining the construct pENTR™/D-TOPO-RBP47b. The CaMV35S promoter of the vectors pGWB642 and pGWB661 was replaced by the pRPS5A promoter, generating the vectors pGWB642(RPS5A) and pGWB661(RPS5A), respectively. pENTR™/D-TOPO-RBP47b was LR-recombined into pGWB642(RPS5A) and pGWB661(RPS5A) to obtain the constructs pGWB642(RPS5A)-RBP47b and pGWB661(RPS5A)-RBP47b, used to generate the lines *YFP-RBP47b* and *TagRFP-RBP47b*, respectively, by transforming the wild type plants using *A. tumefaciens* and the floral dip method.

For the expression of wild type and mutated versions of SMG7, eIF4A1, eIF4A1and eIF4B1 in *Arabidopsis* protoplasts and *Nicotiana tabacum* leaves, we utilized the previously described constructs pGWB442-SMG7, pGWB442-SMG7^K77E^ ^R185E^, pGWB442-SMG7^Δ715-885^, pGWB442-SMG7^Δ917-1010^, and pGWB442-SMG7^Δ702-1059^ (22). The construct pGWB442-SMG7^Δ1-268^ was generated amplifying the cDNA of *SMG7* with the primers smg7_del_14-3-3_topoATG and smg7.TOPO.STOP.R. The fragment was introduced into pENTR™/D-TOPO using blunt-end TOPO® Cloning reactions following LR recombination into pGWB442. The cDNAs of *eIF4A1*, *eIF4A2* and *eIF4B1* were amplified with the following pairs of primers in the respective order: eif4a1.topo.f and eif4a1.nostop.r; EIF4A2.topo.F and eiF4A2.nostop.r; EIF4B1.TOPO.F.real and EIF4B1.NOSTOP.R. The cDNAs were transferred to pENTR™/D-TOPO, and LR recombined into pGWB441, obtaining the constructs pGWB441-eIF4A1, pGWB441-eIF4A2 and pGWB441-eIF4B1. To obtain the constructs The pENTR/D-TOPO-eIF4A1^S146A^, The pENTR/D-TOPO-eIF4A1^T145A^ ^S146A^ and The pENTR/D-TOPO-eIF4A1^S146E^, the pENTR/D-TOPO-eIF4A1 vector was mutagenized with site-directed mutagenesis with the following pairs of primers: eifA1.SDM. S146A.F and eifA1.SDM. S146A.R; eifA1.SDM. T145AS146A.F and eifA1.SDM. T145AS146A.R; eifA1.SDM. S146D.F and eifA1.SDM. S146D.R. Then, LR recombination into pGWB441 was performed to generate the constructs pGWB441-eIF4A1^S146A^, pGWB441-eIF4A1^T145A^ ^S146A^ and pGWB441-eIF4A1^S146E^.

The following constructs for the BiFC experiments were previously described (22): pGW-nY-SMG7, pGW-nY-SMG7^K77E^ ^R185E^, pGW-nY-SMG7^Δ715-885^, pGW-nY-SMG7^Δ917-1010^ and pGW-nY-SMG7^Δ702-1059^. We also generated the constructs pGW-nY-SMG7^Δ1-268^, pGW-eIF4A1-cY, pGW-eIF4A2-cY and pGW-eIF4B1-cY by recombining using LR reaction the entry clones pENTR™/D-TOPO-SMG7 ^Δ1-268^, pENTR™/D-TOPO-eIF4A1, pENTR™/D-TOPO-eIF4A2 and pENTR™/D-TOPO-eIF4B1 with the BiFC vectors pnYGW and pGWcY (66).

### Protein localization and BiFC assay in protoplasts

*Arabidopsis* mesophyll protoplasts were isolated and transfected as described (67). Briefly, mesophyll protoplasts were isolated from leaves of 4-week-old *Arabidopsis* grown on soil in a growth chamber at 22°C under 12 /12 h light/dark cycles. 20-30 leaves were cut using a razor blade and digested in 15 ml of digestion solution (1% cellulase Onozuka R10 [Duchefa], macerozyme R10 [Duchefa], 0,4 M mannitol, 20 mM KCl, 20 mM MES pH=5.7), first for 20 min in vacuum followed by 3 h in dark at room temperature. The released protoplasts were filtered with a 70 µm mesh, washed twice with W5 medium (154 mM NaCl, 125 mM CaCl_2_, 5 mM KCl, 2 mM MES pH=5.7) and stored on ice. For transfection, the protoplasts were resuspended in MMg solution (0,4 mM mannitol, 15 mM MgCL_2_, 4 mM MES pH=5.7) at 3·10^5^ cells/ml. 100 µl of protoplasts were mixed with 15 µg of plasmid and 110 µl of PEG solution (4 g PEG 4000, 2.5 ml mannitol 0.8M, 1 ml CaCl_2_ 1M, 3 ml H_2_0) and incubated 10 min in the dark at RT. 440 µl of W5 medium was added to the mixture and, after centrifugation, the protoplasts were resuspended and incubated in W5 medium for 16 h in the dark at room temperature. Transfected protoplasts were imaged using a Zeiss LSM780 confocal microscope.

### FLIM-FRET assay

Plasmid vectors were transiently expressed in *N. tabacum* (SR1 Petit Havana) leave epidermal cells by the infiltration procedures as described (68). Gene silencing in *Nicotiana tabacum* was suppressed by co-infiltrating the p19 protein from tomato bushy stunt virus cloned into pBIN61 (68). Laser scanning confocal imaging microscope Zeiss LSM 780 AxioObserver equipped with external In Tune laser (488-640 nm, < 3nm width, pulsed at 40 MHz, 1.5 mW) C-Apochromat 63 x water objective, NA 1.2 and, the HPM-100-40 Hybrid Detector from Becker and Hickl GmbH was used for FLIM-FRET data acquisition. FLIM analysis was performed using the Simple-Tau 150N (Compact TCSPC system based on SPC-150N) with DCC-100 detector controller for photon counting. For excitation of YFP and tagRFP, an InTune laser at 490 nm—efficiently matching YFP’s absorption peak while minimizing off-target excitation and phototoxicity— and a 561 nm DPSS lasers were used, respectively (69). The YFP–tagRFP FRET pair was chosen based on adequate spectral overlap between the YFP emission (∼527–530 nm) and the tagRFP excitation (∼555 nm), allowing detection of energy transfer through measurable decreases in YFP fluorescence lifetime (https://www.fpbase.org/spectra/). Although this pair exhibits moderate FRET efficiency, FLIM provides the sensitivity required to detect subtle lifetime shifts indicative of close-range molecular interactions. Zen 2.3 light version from Zeiss was used for processing confocal images. SPCM 64 version 9.8 was used to acquire FLIM data and SPCImage version 7.3 from Becker and Hickl GmbH for data analysis. For each analysis, fluorescence lifetime was collected in range between 500 and 5000 ps (A: below 500 ps = background noise, chloroplasts and dirt/dust, B: above 5000 ps = unspecific signal caused by i.e. P-body movement, autofluorescent foci/other cellular compartments) using 2D correlation. A multiexponential decay model was used for fitting. Student’s t-test was done to evaluate the significant difference in the average lifetime between different groups.

### Cytology

Pollen count and viability were determined by Alexander staining as described (70), and imaged using the transmitted light microscope Zeiss Axioscope.A1(objective 20x/0.5), with the Axiocam 105 camera and the software Visitrone Visiview. Meiosis was assessed by DAPI staining of PMCs in whole anthers as described (35), and imaged on a Zeiss LSM780 confocal microscope.

### Root sample preparation for protein localization

Protein localization in *Arabidopsis* roots was performed using roots of 4 days old seedlings. Seedlings were fixed in 4% formaldehyde in FB (1 mM EDTA, 0.1% Triton X-100 in PBS 1X pH7). The sample was incubated for 1 h within vacuum, followed by 3 washes with FB. The sample was stained with 1:1000 dilution of SR2200 in PB for 30 min and washed 3 times with FB. Finally, the sample was placed in a microscope slide with VECTASHIELD® Antifade Mounting Media for further imaging.

### Anther sample preparation for protein localization

Protein localization in *Arabidopsis* anthers was performed as described in Capitao et al., 2021.^3^ Inflorescences were fixed by vacuum incubation with 4% formaldehyde in FB for 15 min, followed by 45 min of incubation without vacuum. The sample was washed 3 times with FB before dissecting the meiotic floral buds to isolate meiotic anthers, and then they were incubated 1 hour with 5 µg/ml DAPI in FB. Next, the anthers were washed 3 times with FB, 1 time with FB at 60°C for 10 min, 1 time with FB on ice, and 1 time with FB at room temperature. Finally, the anthers were placed in a microscope slide with VECTASHIELD® Antifade Mounting Media to be subsequently imaged.

### Laser scanning confocal imaging

Laser scanning confocal imaging was performed in two systems. The Inverted microscope Zeiss Axio Observer.Z1 with confocal unit LSM 780 was used to image protoplasts, roots (quantification) and anthers. For the protoplasts we used the objective LCI Plan-Neofluar 63x/1.3 1mm Korr DIC M27. For the roots and anthers, we used the objective C-Apochromat 63x/1.2 W Korr UV-VIR-IR M27. The Inverted microscope Zeiss Axio Observer.7 with confocal unit LSM 880 was used for the colocalization imaging in roots. In this case, we used the objective Plan-Apochromat 63x/1.4 Oil DIC M27.

### Super-resolution imaging

To obtain a resolution beyond the Abbe diffraction limit of the conventional light microscope we used the motorized inverted microscope ZEISS Elyra 7 with lattice illumination pattern for 3D structured illumination (Lattice SIM). This system was used to image RNP granules of roots and meiocytes. We used the objective Plan-Apochromat 63x/1.4 Oil DIC M27 and the 2 x PCO edge sCMOS camera, 1280 x 1280, pixel size 6.5 μm × 6.5 μm. For excitation we used 488 nm and 561nm lasers. The raw images were acquired using a dimension of 1024 x 1024 pixels (pixel size 63 nm) and 9 phases.

### Live cell imaging

Live cell imaging of *Arabidopsis* pollen mother cells was performed using Light sheet microscopy as previously described (71). Anthers were imaged in Light sheet Z.1 microscope (Zeiss) (Objective W Plant-Apochromat 20x/1.0 DIC) in 15 min time increments.

### Image processing

Images obtained by laser scanning confocal imaging were processed for image presentation with the software ZEN (blue and black edition). The 3D segmentation and the signal quantification of RNP granules from roots and meiocytes was performed with the Imaris 10.2.0 microscopy image analysis software, utilizing the volume rendering. The images obtained with the super-resolution microscope ZEISS Elyra 7 were processed in ZEN Black 3.0 SR (Zeiss), utilizing the 3D SIM2 algorithm, with output sampling = 2 and scaled to raw. The software Imaris 10.2.0 was used for 3D segmentation and volume rendering.

### Western blot analysis

Protein extracts were prepared by grinding leaves, stems and inflorescences from 35-day old *SMG7-MYC* line and wild type control in liquid nitrogen. Whole protein extract was obtained after 15 min incubation at 4°C with protein extraction buffer (50 mM Tris-HCl pH 7.5, 150 mM NaCl, 0.1% Nonidet P-40, 10% Glycerol, 1 mM PMSF) supplemented with protease inhibitor cocktail (Roche). After 15 min of incubation at 4°C, the sample was centrifugated at 17500 rpm. The protein extract contained in the supernatant fraction was quantified, and 45 µg of protein were loaded onto an SDS-PAGE gel. For Western blotting, SMG7-MYC was detected using anti-MYC polyclonal antibody (ab9106 Abcam).

### Immunoprecipitation and Mass Spectrometry

Leaves, stems and inflorescences from 35-day old *SMG7-MYC* line and wild type control were grinded in liquid nitrogen and processed using the EZview™ Red Anti-c-Myc Affinity Gel (Merck). Briefly, 2 g of grinded tissue was mixed with 5 ml of protein extraction buffer (50 mM Tris-HCl pH 7.5, 150 mM NaCl, 0.1% Nonidet P-40, 10% Glycerol, 1 mM PMSF) supplemented with protease inhibitor cocktail (Roche) and PhosStop phosphatase inhibitor cocktail (Roche). After 15 min of incubation at 4°C, the sample was centrifugated at 17500 rpm. 10 mg of protein were incubated with 50 µl of affinity gel for 1 h 30 min with rotation. Finally, the sample was centrifugated for 8200 g and washed 5 times with extraction buffer.

In-gel digestion: The immunoprecipitated proteins were separated using 1D-SDS-PAGE gel electrophoresis (12% gel, one replicate per gel) and whole gel lanes selected for analysis were excised in the form of bands (30 bands per lane). After destaining and washing procedures, each band was washed by 50% ACN/NaHCO_3_ and pure ACN and the gel pieces were incubated with 125 ng trypsin (sequencing grade; Promega) in 50mM NaHCO_3_. The digestion was performed for 2 h at 40 °C on a Thermomixer (750 rpm; Eppendorf). Tryptic peptides were extracted into LC-MS vials by 2.5% formic acid (FA) in 50% ACN with addition of polyethylene glycol (20,000; final concentration 0.001%) and concentrated in a SpeedVac concentrator (Thermo Fisher Scientific).

LC-MS/MS analyses of individual gel band digests were done Ultimate 3000 RSLCnano system connected to Orbitrap Elite hybrid spectrometer (Thermo Fisher Scientific). Prior to LC separation, tryptic digests were online concentrated and desalted using trapping column (100 μm × 30 mm) filled with 3.5-μm X-Bridge BEH 130 C18 sorbent (Waters). After washing of trapping column with 0.1% FA, the peptides were eluted (flow 300 nl/min) from the trapping column onto a Acclaim Pepmap100 C18 column (2 µm particles, 75 μm × 250 mm; Thermo Fisher Scientific) by the following gradient program (mobile phase A: 0.1% FA in water; mobile phase B: 80% ACN containing 0.1% FA): the gradient elution started at 1% of mobile phase B and increased from 1% to 56% during the first 50 min (1% in the 1st, 30% in the 30th and 56% in 50th min), then increased linearly to 80% of mobile phase B in the next 5 min and remained at this state for the next 10 min. Equilibration of the trapping column and the column was done prior to sample injection to sample loop. The analytical column outlet was directly connected to the Nanospray Flex Ion Source (Thermo Fisher Scientific).

MS data were acquired in a data-dependent strategy selecting up to top 10 precursors based on precursor abundance in the survey scan (350-2000 m/z). The resolution of the survey scan was 60,000 (at m/z 400) with a target value of 1×10^6^ ions, one microscan and maximum injection time of 200 ms. HCD MS/MS spectra (relative fragmentation energy of 32 %, resolution 15,000 at m/z 400) were acquired with a target value of 50 000 ions with m/z range adjusted according to actual precursor mass and charge. The maximum injection time for MS/MS was 500 ms. Dynamic exclusion was enabled for 45 s after one MS/MS spectra acquisition and early expiration was disabled. The isolation window for MS/MS precursor isolation was set to 2 m/z.

For data evaluation, MaxQuant software (v1.6.17.0) (72) with inbuild Andromeda search engine (73) was used. Search was done against protein databases of *Arabidopsis thaliana* (27,469 protein sequences, version from 2021-11-17, downloaded from: https://ftp.uniprot.org/pub/databases/uniprot/current_release/knowledgebase/reference_proteom es/Eukaryota/UP000006548/UP000006548_3702.fasta.gz) and cRAP contaminants (112 sequences, version from 2018-11-22, downloaded from http://www.thegpm.org/crap). The following variable modifications were set for the database search: oxidation (M), propionamide (C), and acetylation (Protein N-term). Enzyme specificity was tryptic with two permissible miscleavages. Only peptides and proteins with false discovery rate threshold under 0.01 were considered. Relative protein abundance was assessed using protein intensities calculated by MaxQuant. Match between runs was enabled and set using the fractions scheme specifying individual bands as different consecutive fractions (1–30) using the same fraction number for the corresponding gel area from the two samples of one replicate but different fraction number between replicates.

Intensities of reported proteins were further evaluated using software container environment (https://github.com/OmicsWorkflows/KNIME_docker_vnc; version 4.1.3a). Processing workflow is available upon request and it covered, in short, reverse hits and contaminant protein groups (cRAP) removal, protein group intensities log2 transformation and normalization (loessF). Replicate-wise ratios were calculated using the normalized data and evaluated to filter for putative proteins candidates list. A complete list of proteins within each protein group can be viewed in the supporting material (Table S1). Mass spectrometry proteomics data were deposited to the ProteomeXchange Consortium via PRIDE (74) partner repository under dataset identifier PXD061406 and 10.6019/PXD061406

### Protein co-immunoprecipitation and western

Leaves and inflorescences from 35-day old *SMG7-MYC*, *eIF4A1-YFP* and *SMG7-MYC eIF4A1-YFP* lines were grinded in liquid nitrogen. Whole protein extract was obtained after 15 min incubation at 4°C with protein extraction buffer (50 mM Tris-HCl pH 7.5, 150 mM NaCl, 0.1% Nonidet P-40, 10% Glycerol, 1 mM PMSF) supplemented with protease inhibitor cocktail (Roche). After 15 min of incubation at 4°C, the sample was centrifugated at 17500 rpm. The protein extract contained in the supernatant fraction was quantified, and 2.5 mg of protein were incubated with GFP-Trap® Magnetic Agarose (AB_2631358 ChromoTek) for 3 h 30 min at 4°C with rotation. After 4 washes with extraction buffer, proteins were eluted from the affinity beads by adding 4x Leammli sample buffer. For Western blotting, proteins were detected using anti GFP monoclonal antibody (3H9) (AB_10773374 ChromoTek) and anti-MYC polyclonal antibody (ab9106 Abcam).

### Protein structure model building

The Alphafold-3 model of the SMG7-eIF4A1 interaction was downloaded from the Alphafold Server (37) and visualized and prepared for presentation with UCSF ChimeraX (75).

### Quantification and statistical analysis

Quantification analysis of condensates was performed using Imaris 10.2.0 software. Counting of viable pollen grains was performed using the Count tool of Adobe Photoshop 13.0 CS6 Extended.

Data analysis and graph constructions were performed using GraphPad Prism 7 software. Significance testing was conducted with Student’s t tests with a significance threshold of p < 0.05, and One-way ANOVA followed by Tukey’s post-hoc test. Further details can be found in the figure legends.

## Acknowledgements

This work was supported by the Czech Science Foundation (EXPRO grant 23-07969X). We further acknowledge the core facilities of CEITEC Masaryk University for support in obtaining and analysing scientific data: CELLIM funded by MEYS CR (LM2023050 Czech-BioImaging), Proteomics Core Facility of CIISB, Instruct-CZ Centre, supported by MEYS CR (LM2023042, CZ.02.01.01/00/23_ 015/0008175, e-INFRA CZ (ID:90254), and Plant Science Core Facility.

