## Supplementary material for "SMG7 and eIF4A constitute a homeostatic module controlling P-body condensation and function of Meiotic bodies": Suplemental information

Table S1 and S2

Figures S1-S7

| Accession | Locus | Description | Coverage | Number of Peptides (Unique) | Fold Change (Replica 1) | Fold Change (Replica 2) | Fold Change (Replica 3) |
|---|---|---|---|---|---|---|---|
| A9QM73 | At5G19400 | SMG7 | 0,50 | 46(46) | qual | qual | qual |
| P41377 | At1G54270 | eIF4A2 - Eukaryotic translation initiation factor 4A-2 | 0,63 | 28(1) | 2,93 | 3,10 | 2,77 |
| F4JEL5 | AT3G13920 | eIF4A1 - Eukaryotic translation initiation factor 4A-1 | 0,77 | 33(9) | 2,93 | 3,72 | 2,80 |
| P21218 | AT4G27440 | PORB - Protochlorophyllide reductase B | 0,50 | 19(15) | 2,67 | 4,80 | 2,67 |
| Q944S1 | AT1G59990 | RH22 - DEAD-box ATP-dependent RNA helicase 22 | 0,32 | 17(17) | 3,94 | 2,87 | 4,29 |
| Q948R9 | AT3G17170 | RFC3 - Regulator of fatty acid composition 3 | 0,38 | 10(10) | qual | 3,19 | 2,64 |
| F4I3P9 | AT1G63680 | MURE | 0,35 | 21(21) | 7,19 | qual | 3,80 |
| Q9LPG6 | AT1G53500 | RHM2 | 0,33 | 20(7) | 2,62 | 7,31 | 4,16 |
| Q9LH76 | AT3G14790 | RHM3 | 0,32 | 19(7) | 2,70 | 3,57 | 4,31 |
| P92948 | At1G09770 | CDC5 - Cell division cycle 5-like protein | 0,19 | 13(13) | 2,57 | 2,75 | qual |
| F4ICK7 | AT1G32130 | IWS1 - Transcription elongation factor (TFIIS) family protein | 0,26 | 11(11) | qual | qual | qual |
| F4KFP7 | At5G24060 | Pentatricopeptide repeat-containing protein-like protein | 0,19 | 8(8) | qual | qual | qual |
| B3H778 | At4G24830 | Argininosuccinate synthase | 0,09 | 3(3) | qual | qual | qual |
| Q9M0A7 | AT4G30530 | GGP1 - Gamma-glutamyl peptidase 1 | 0,18 | 4(4) | qual | qual | qual |
| Q9C522 | At3G06650 | ACLB1 - ATP-citrate lyase B-1 | 0,07 | 4(4) | qual | qual | qual |
| B3H757 | AT4G29810 | MKK2 - Mitogen-activated protein kinase kinase 2 | 0,09 | 3(3) | qual | qual | qual |
| Q39048 | At4G24510 | ECERIFERUM 2 | 0,05 | 2(2) | qual | qual | qual |
| Q8LEF3 | At5G46030 | ELF1 - Transcription elongation factor 1 homolog | 0,30 | 1(1) | qual | qual | qual |
| Q9LXN4 | At3G44530 | HIRA | 0,02 | 1(1) | 4,59 | qual | qual |

### Table S1. Mass spectrometry analysis of the SMG7 interactome, related to Figure 1

List of proteins identified by LC–MS in pull-downs from wild type and SMG7:MYC line. The list includes the proteins with ≥2.5 enrichment (FC) in the three independent immunoprecipitation experiments. qual = qualitative enriched (only present in the SMG7:MYC sample).

| PRIMER | SEQUENCE (5’->3’) |
|---|---|
| UBP1b.BamHI.V2.F | TTATGATATCTATGCAGAGGTTGAAGCAGCAGC |
| UBP1b.6.R | GGGGGAGATTTCCACTTGGGATTGG |
| eIF4A1.Prom.F | CACCCTGCGTCAGCCGATCCGAATTCG |
| eif4A1.nostop.r | CAGCAGATCGGCCACGTTCGAAGGC |
| eif4A2.Promoter.Topo.F | CACCAAGCCAACACTCACCCTGCGTCC |
| eiF4A2.nostop.r | CAGCAAATCAGCCACGTTTGAGGG |
| DCP1_FP | CACCTGGTCTTAACAAGGCAGGCA |
| DCP1_RP | TTGTTGAAGTGCATTTTGTAAAGTTCGGGTA |
| UBPb1.EcoRI.F.V1 | GGAAACAAAATGCGATGC |
| UBPb1EcoRV.R.V2 | CTGGTAGTACATGAGCTGCTGCGCG |
| RBP47b_cacc_topo_F_V2 | CACCATGCAGACAACCAACGGCTC |
| RBP47b_stop_R | TCAATTCTCCCCATGATAGTTGTTGG |
| smg7_del_14-3-3_topoATG | CACCATGTATGAAAAATTGTTTGTGCCCTCC |
| smg7.TOPO.STOP.R | TCACACAAAGTGACGACTCGACC |
| EIF4A1.topo.F | CACCATGGCAGGATCTGCACCAGAAGGC |
| EIF4A2.topo.F | CACCATGGCAGGATCCGCACCGGAAGGAAC |
| EIF4B1.TOPO.F.real | CACCATGTCGAAAGCTTGGGGTGGAATTGG |
| EIF4B1.NOSTOP.R | CCATCCTTCCCTAGAGGAAG |

**Table S2. List of primers**

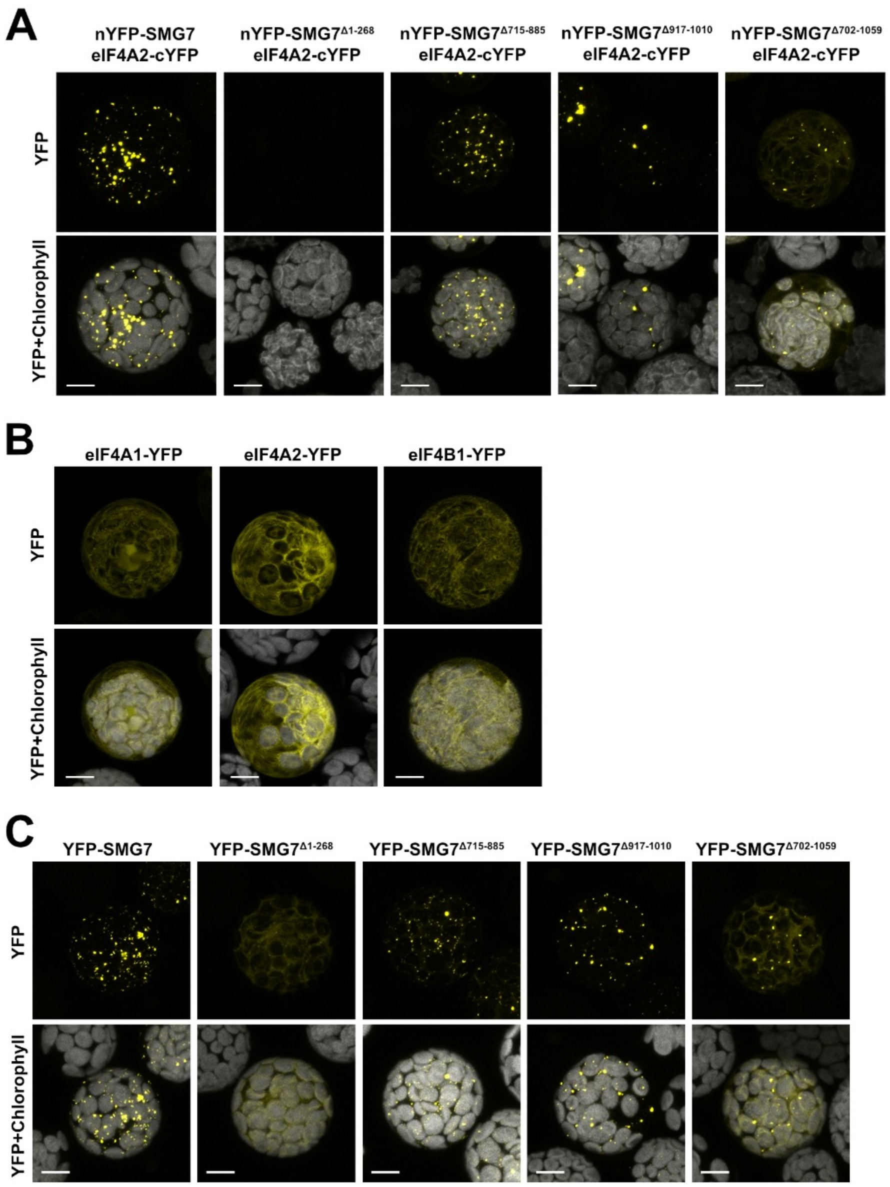


### Figure S1. BiFC and localization of eIF4A1/2 and SMG7 in mesophyll protoplasts

**(A)** BiFC assay in mesophyll protoplasts for eIF4A2 interaction with the truncated versions of SMG7. Scale bar = 10 μm. **(B)** Mesophyll protoplasts transiently transfected with the eIF4A1-YFP, eIF4A2-YFP and eIF4B1-YFP constructs. Scale bar = 10 μm. **(C)** Mesophyll protoplasts transiently transfected with the YFP-SMG7 construct and its mutated derivatives. Scale bar = 10 μm.

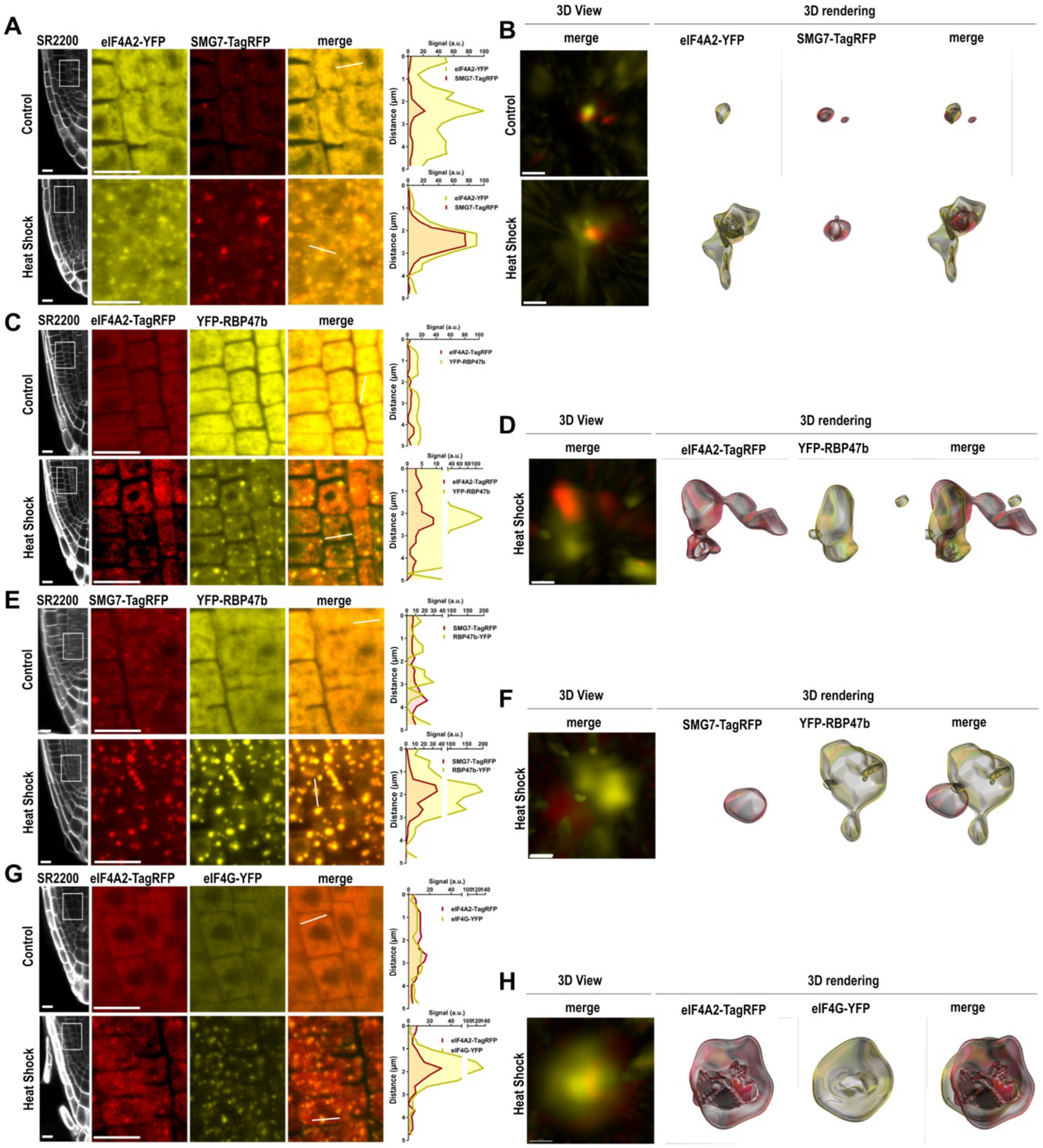


### Figure S2. eIF4A2 localizes to P-bodies and SGs in *Arabidopsis* roots

**(A),(C),(E),(G),(I)** Confocal micrographs of root cells co-expressing indicated combinations of reporter proteins. Counterstaining with SR2200 dye was used to visualize cell walls. Diagrams on the left show superimposed intensity profiles of YFP and TagRFP signals measured along the lines indicated in the corresponding micrographs. Scale bar = 10 µm. **(B),(D),(F),(H),(J)** Super-resolution micrographs of indicated protein condensates visualized by 3D view and 3D rendering using Imaris software. Scale bar = 0.5 µm.

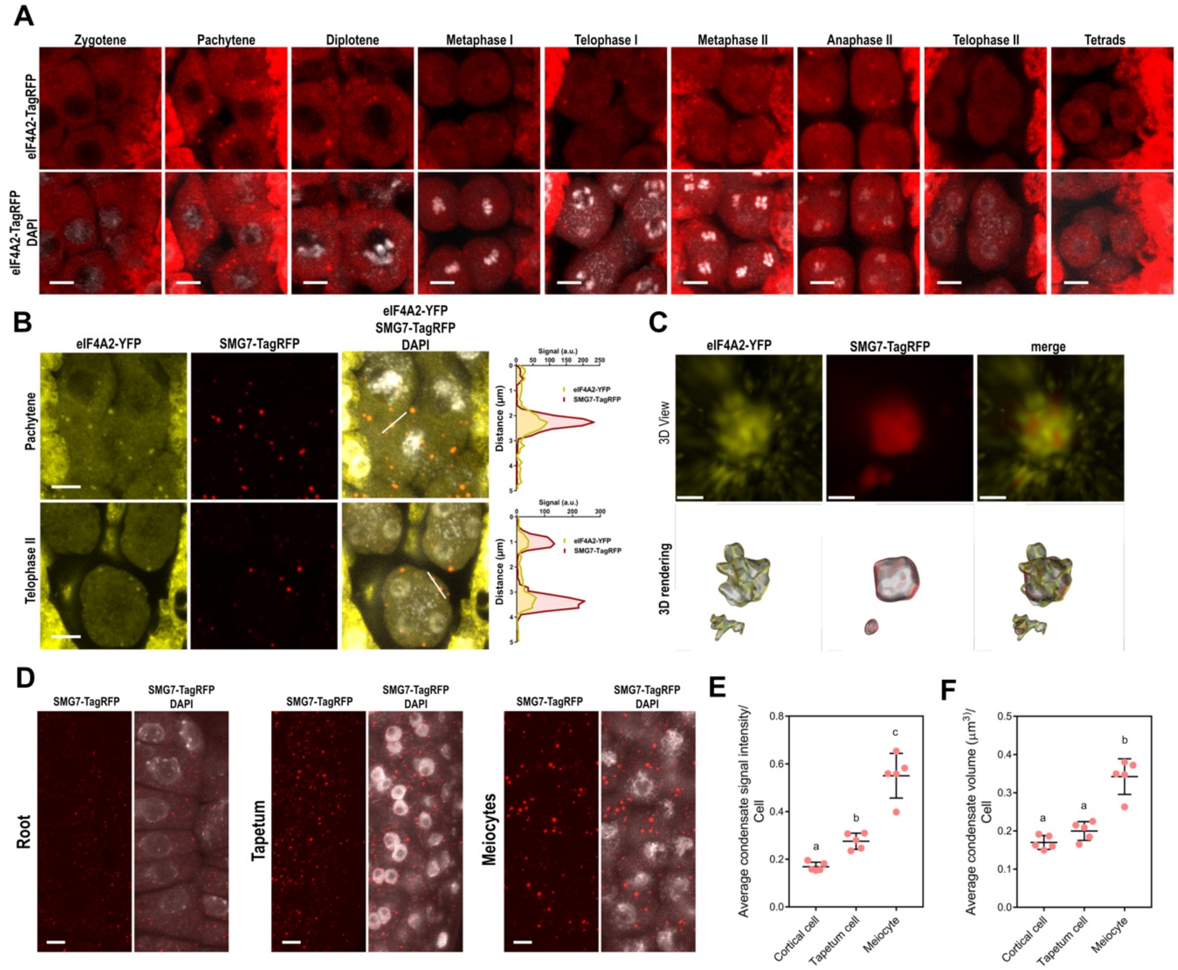


### Figure S3. eIF4A2 localization during meiosis

**(A)** Confocal micrographs of *Arabidopsis* anthers showing localization of eIF4A2-TagRFP in the course of meiosis. DNA is counterstained with DAPI. Scale bars = 5 µm. **(B)** Confocal micrographs of *Arabidopsis* meiocytes co-expressing eIF4A2-TagRFP and SMG7-TagRFP. Diagrams on the left show superimposed intensity profiles of YFP and TagRFP signals measured along the lines indicated in the corresponding micrographs. Scale bar = 10 µm. **(C)** Super-resolution micrographs of indicated protein condensates visualized by 3D view and 3D rendering using Imaris software. Scale bar = 0.5 µm. **(D)** Confocal micrographs of root cortical cells, tapetal cells, and pollen mother cells expressing *SMG7-TagRFP.* Scale bar = 5 µm. Dot plots showing **(E)** average of the signal intensity and **(F)** average volume of the condensates from (A-C) (mean, SD, n = 5, One-way ANOVA followed by Tukey's post-hoc test $p < 0.05$).

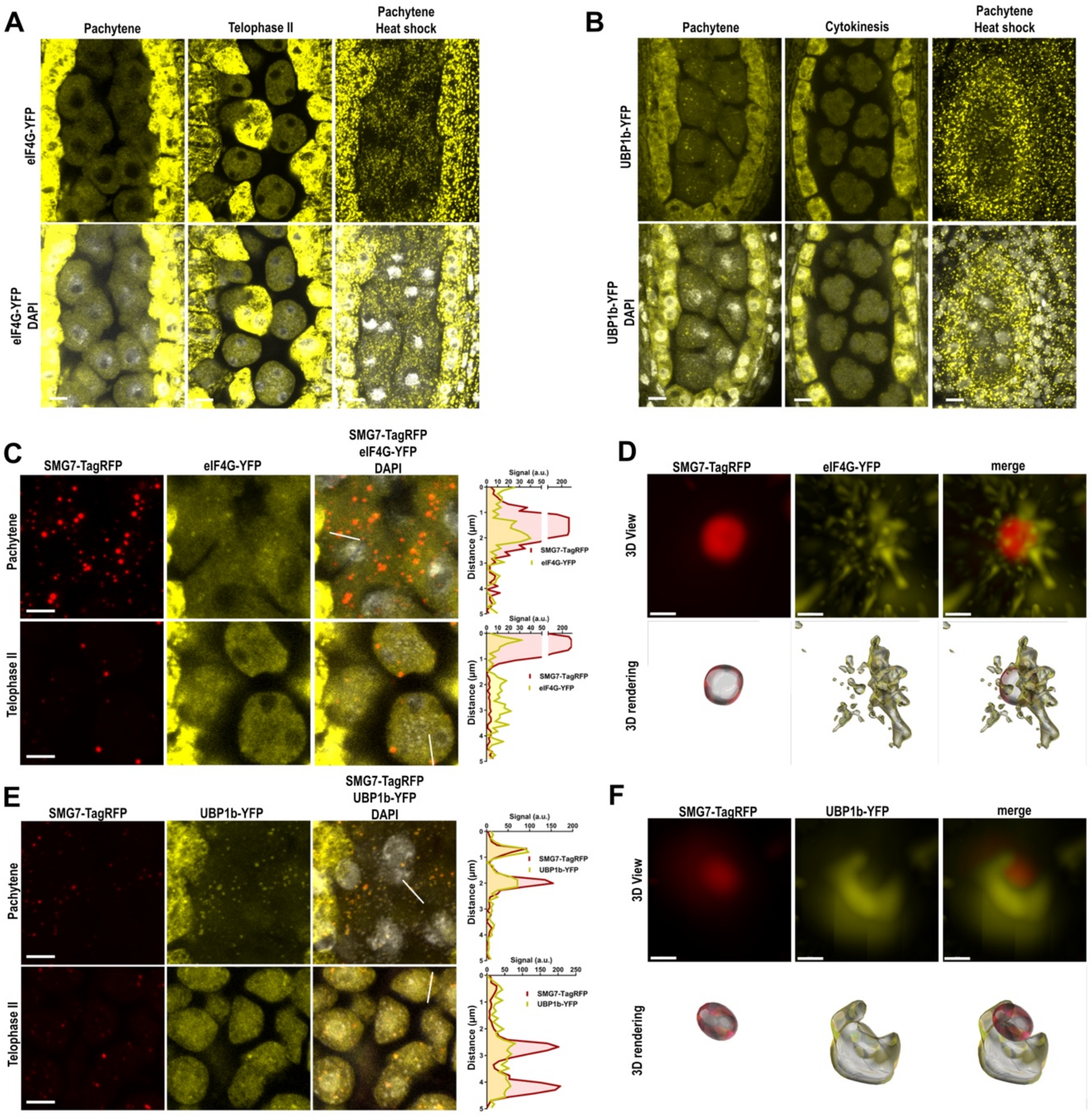


**Figure S4. Composition of M-bodies.**

**(A)** Confocal micrographs of anther lobes showing expression and localization of eIF4G-YFP in meiocytes and the surrounding tapetum upon heat shock or under control conditions without heat shock application. Scale bar = 5 µm. **(B)** Confocal micrographs of anther lobes showing expression and localization of UBP1b-YFP in meiocytes and the surrounding tapetum upon heat shock or under control conditions without heat shock application. Scale bar = 5 µm. **(C)** Confocal micrographs of *Arabidopsis* meiocytes co-expressing SMG7-TagRFP and eIF4A-YFP. Diagrams on the left show superimposed intensity profiles of YFP and TagRFP signals measured along the lines indicated in the corresponding micrographs. Scale bar = 5 µm. **(D)** Super-resolution micrographs of SMG7-TagRFP / eIF4A-YFP condensates visualized by 3D view and 3D rendering using Imaris software. Scale bar = 0.5 µm. **(E)** Confocal micrographs of *Arabidopsis* meiocytes co-expressing SMG7-TagRFP and UBP1b-YFP. Diagrams on the left show superimposed intensity profiles of YFP and TagRFP signals measured along the lines indicated in the corresponding micrographs. Scale bar = 5 µm. **(F)** Super-resolution micrographs of SMG7-TagRFP / UBP1b-YFP condensates visualized by 3D view and 3D rendering using Imaris software. Scale bar = 0.5 µm.

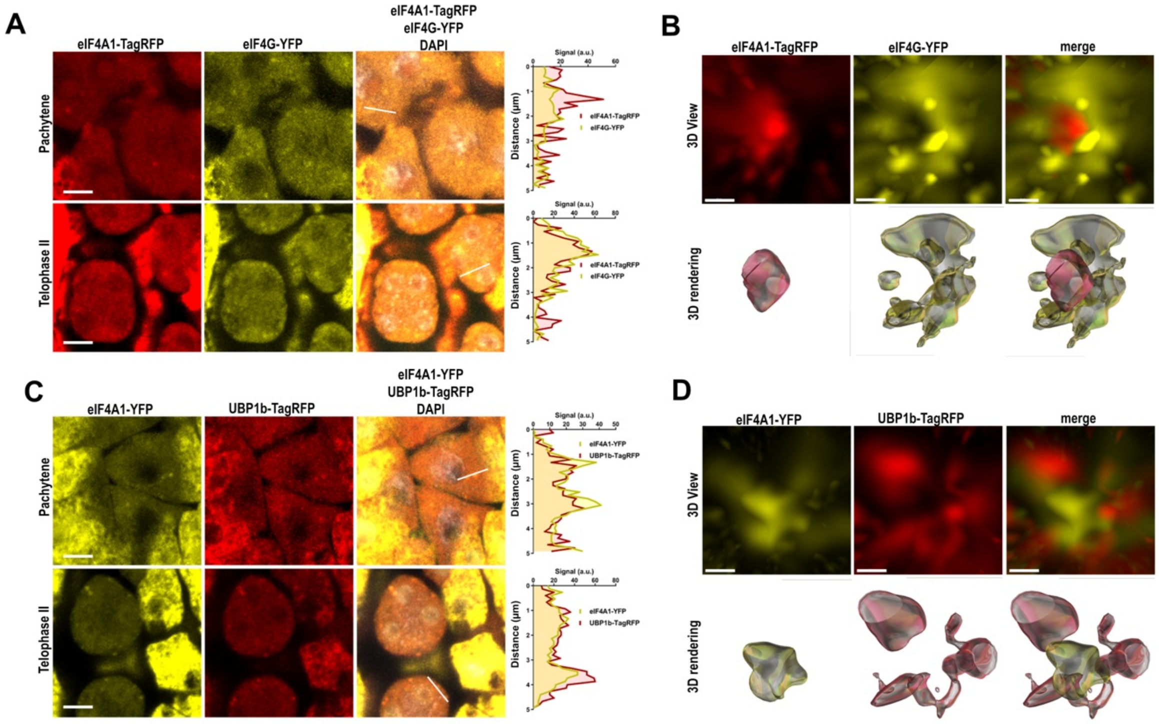


**Figure S5. Co-localization of eIF4A1 with eIF4G and UBP1b in M-bodies**

**(A)** Confocal micrographs of *Arabidopsis* meiocytes co-expressing eIF4A1-TagRFP and eIF4G-YFP. Diagrams on the left show superimposed intensity profiles of YFP and TagRFP signals measured along the lines indicated in the corresponding micrographs. Scale bar = 5 µm. **(B)** Super-resolution micrographs of eIF4A1-TagRFP / eIF4G-YFP condensates visualized by 3D view and 3D rendering using Imaris software. Scale bar = 0.5 µm. **(C)** Confocal micrographs of *Arabidopsis* meiocytes co-expressing eIF4A1-YFP and UBP1b-TagRFP. Diagrams on the left show superimposed intensity profiles of YFP and TagRFP signals measured along the lines indicated in the corresponding micrographs. Scale bar = 5 µm. **(D)** Super-resolution micrographs of eIF4A1-YFP / UBP1b-TagRFP condensates visualized by 3D view and 3D rendering using Imaris software. Scale bar = 0.5 µm.

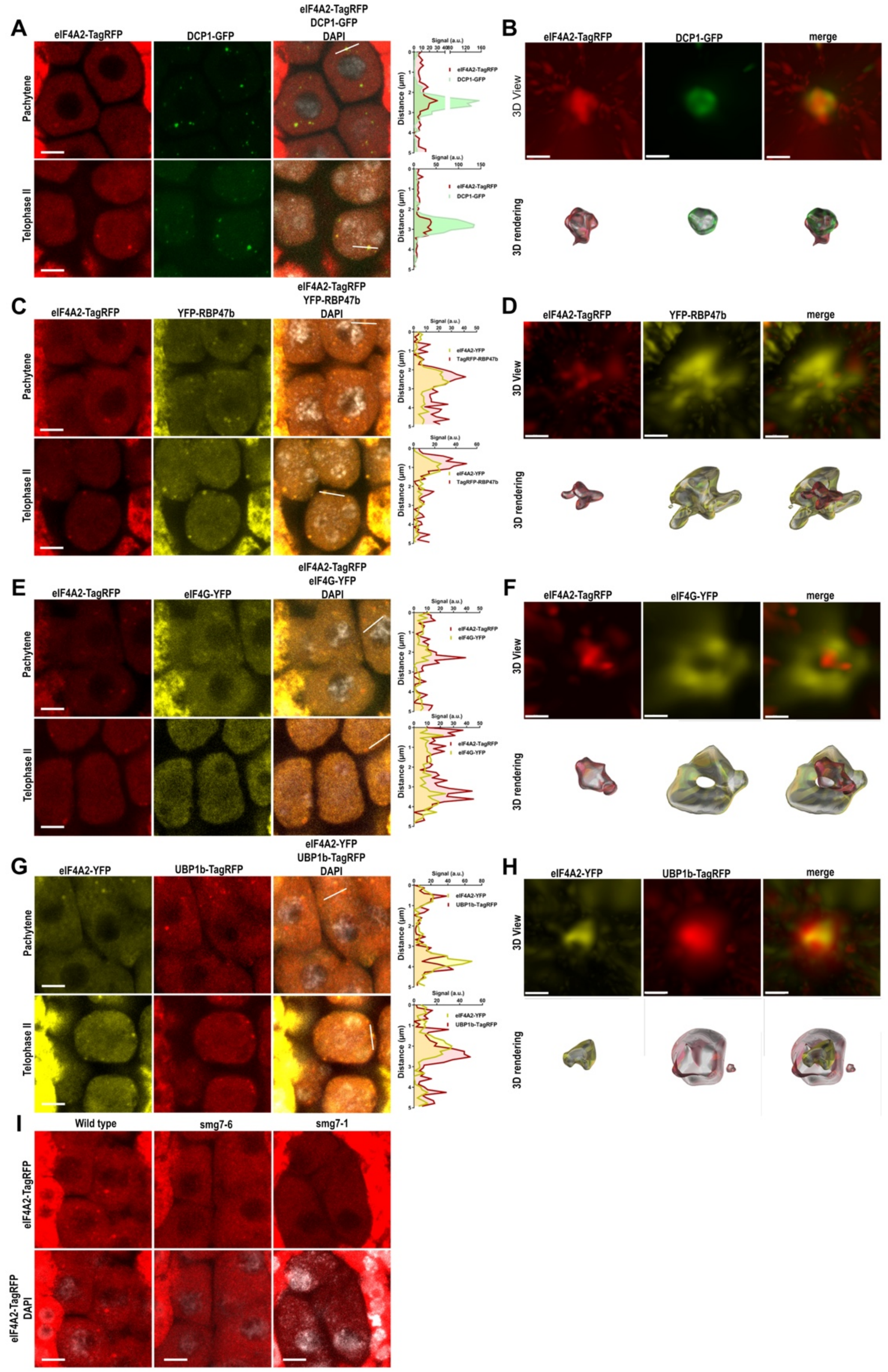


**Figure S6. Localization of eIF4A2 in M-bodies.**

**(A),(C),(E),(G)** Confocal micrographs of meiocytes co-expressing indicated combinations of reporter proteins Diagrams on the left show superimposed intensity profiles of YFP and TagRFP signals measured along the lines indicated in the corresponding micrographs. Scale bar = 10 μm. **(B),(D),(F),(H)** Super-resolution micrographs of indicated protein condensates visualized by 3D view and 3D rendering using Imaris software. Scale bar = 0.5 μm. **(I)** Confocal micrographs of *Arabidopsis* meiocytes of wild type, *smg7-6* and *smg7-1* plants expressing eIF4A2-TagRFP. Scale bars, 5 μm.

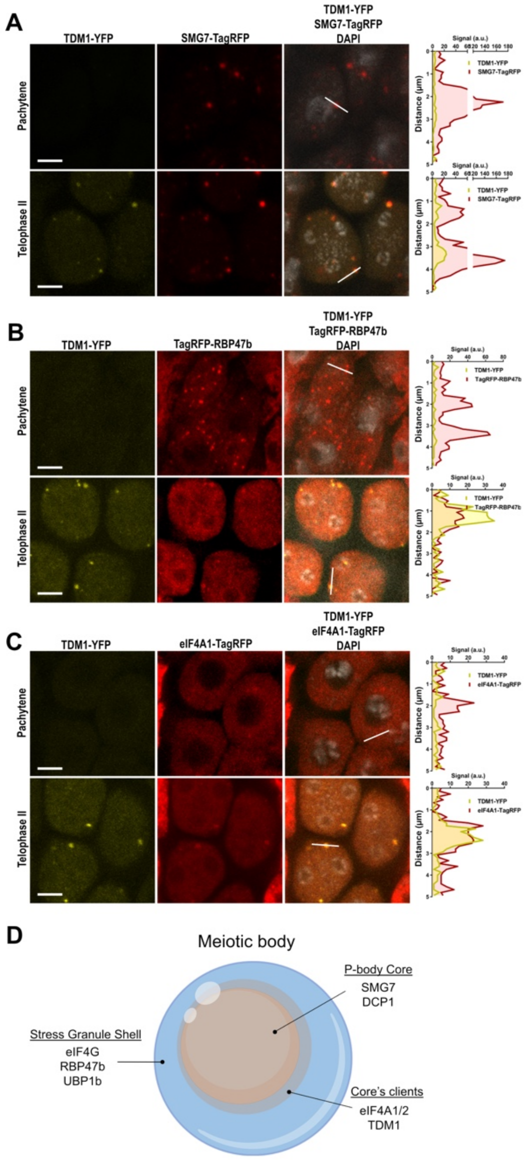


### Figure S7. Localization of TDM1 in M-bodies

**(A), (B), (C)** Confocal micrographs of meiocytes co-expressing TDM1-YFP with indicated combinations of reporter proteins. Diagrams on the left show superimposed intensity profiles of YFP and TagRFP signals measured along the lines indicated in the corresponding micrographs. Scale bar = 10 µm. **(D)** Graphical representation of the M-body structure with indicated proteins.
